# A Notch and Su(H) dependent enhancer complex coordinates expression of *nab* in *Drosophila*

**DOI:** 10.1101/036038

**Authors:** Elizabeth Stroebele, Albert Erives

## Abstract

The transcription factor Suppressor of Hairless and its co-activator, the Notch intracellular domain, are polyglutamine (pQ)-rich factors that target enhancer elements and interact with other locally-bound pQ-rich factors. To understand the functional repertoire of such enhancers, we identify conserved regulatory belts with binding sites for the pQ-rich effectors of both Notch and BMP/Dpp signaling, and the pQ-deficient tissue selectors Apterous (Ap), Scalloped (Sd), and Vestigial (Vg). We find that the densest such binding site cluster in the genome is located in the BMP-inducible *nab* locus, a homolog of the vertebrate transcriptional co-factors *NAB1*/*NAB2*. We report three major findings. First, we find that this *nab* regulatory belt is a novel enhancer driving dorsal wing margin expression in regions of peak phosphorylated-Mad in wing imaginal discs. Second, we show that Ap is developmentally required to license the *nab* dorsal wing margin enhancer (DWME) to read-out Notch signaling in the dorsal wing compartment. Third, we find that the *nab* DWME is embedded in a complex of intronic enhancers, including a wing quadrant enhancer, a proximal wing disc enhancer, and a larval brain enhancer. This enhancer complex coordinates global *nab* expression via both tissue-specific activation and inter-enhancer silencing. We suggest that DWME integration of BMP signaling maintains *nab* expression in proliferating margin descendants that have divided away from Notch-Delta boundary signaling. As such, uniform expression of genes like *nab* and *vestigial* in proliferating compartments would typically require both boundary and non-boundary lineage-specific enhancers.

---

Notch signaling is a key animal innovation used to ensure different cell fates in adjacent cells (Artavanis-Tsakonas 1999; Barad *et al*. 2011; Guruharsha *et al*. 2012). Cell-cell signaling between membrane-bound Notch receptor and its membrane-bound ligands, Delta and Serrate/Jagged, leads to cleavage and nuclear import of the Notch intracellular domain (NICD) (Schroeter *et al*. 1998). In the nucleus, NICD binds the transcription factor Suppressor of Hairless, Su(H), to induce or permit activation of target genes via their transcriptional enhancers (Fortini and Artavanis-Tsakonas 1994). This role of Su(H) is further complexified because it can recruit the repressor Hairless in the absence of NICD, further promoting divergent cell fate trajectories (Bang *et al*. 1995; Barolo *et al*. 2002; Maier *et al*. 2011; Ozdemir *et al*. 2014). This simple operation is central to diverse developmental contexts: **(*i*)** tissue compartment boundaries and organizers (signaling across a linear border defining two domains in an epithelium), **(*ii*)** proneural clusters (signaling across a circular border encircling a field of cells in an epithelium), **(*iii*)** somatic germline niches (signaling between germline stem cells and neighboring somatic cells), **(*iv*)** neural stem progenitors and sensory organ precursors (signaling between one cell and surrounding epithelial cells), and **(*v*)** asymmetric cell fate lineages (signaling between two dividing daughter cells). For this reason, Notch signaling to a tissue-specific transcriptional enhancer is frequently integrated with different, context-specific, signaling cues throughout development (Voas and Rebay 2004; Ward *et al*. 2006; Liu and Posakony 2012; Housden *et al*. 2014).

Notch-target enhancers can be characterized as either Notch instructive or Notch permissive (Bray and Furriols 2001), although other types are also evident (Janody and Treisman 2011). Examples of both instructive and permissive Notch-target enhancers are known for genes expressed in the embryo and imaginal discs of *Drosophila*. In the early embryo, the *E(spl)m8* enhancer is a neurogenic target of an instructive Notch signal (Furukawa *et al*. 1995; Lecourtois and Schweisguth 1995; Schweisguth 1995). Misexpression of NICD in an ectopic stripe spanning the entire dorsal–ventral (D–V) axis drives over-expression of *E(spl)m8* throughout the D–V axis, except in the mesoderm where it is repressed by the C2H2 zinc finger repressor Snail (Cowden and Levine 2002). In contrast, the Notch-target, Su(H)-dependent, *sim* mesoectodermal enhancer and *rhomboid* (*rho*) neurogenic ectoderm enhancer (NEE) are driven ectopically in a way that is limited by the nuclear morphogenic gradient of Dorsal, which also targets these enhancers (Cowden and Levine 2002; Markstein *et al*. 2004; Crocker *et al*. 2010). Thus, in the context of the *sim* and *rho* enhancers, the Notch signal is only permissive because it is not sufficient for expression.

Wing margin enhancers drive expression along the border separating dorsal and ventral compartments of wing imaginal discs (Jack *et al*. 1991; Williams *et al*. 1994; Lecourtois and Schweisguth 1995; Neumann and Cohen 1996b). Wing margin enhancers from *E(spl)m8* and *cut* use Notch signaling instructively, whereas enhancers from *vestigial* (*vg*) and *wingless* (*wg*) use the signal permissively (Janody and Treisman 2011). Tellingly, these enhancer types respond differently to mutations of genes encoding subunits of the Mediator co-activator complex such as Med12 and Med13, which are required for Notch signaling (Janody and Treisman 2011). In clones deleted for Med12 and Med13, Notch-instructive margin/boundary enhancers from *E(spl)m8* and *cut* fail to drive any expression, while Notch-permissive boundary/margin enhancers from *vestigial* (*vg*) and *wingless* (*wg*) drive expression that is limited to cells close to the margin. Thus, diverse developmental enhancers encode contextual information specifying whether the Notch signal is sufficient and therefore instructive, or only permissive of activation by other integral patterning signals.

How an enhancer integrates Notch signaling with other signaling pathways in a developmental context is an important question. Some insight comes from studies of the nonhomologous, Notch-permissive NEEs at *rho*, *vn*, *brk*, *vnd*, and *sog* (Erives and Levine 2004; Crocker *et al*. 2008, 2010; Crocker and Erives 2013; Brittain *et al*. 2014). These enhancers are driven by the Dorsal/Rel morphogenic gradient patterning system of *Drosophila*. Activation is mediated by a pair of linked binding sites for Dorsal and Twist:Daughterless (Twi:Da) heterodimers, as well as by a separate site for the MADF BESS-domain containing factor Dip3, which is important for SUMOylation of Dorsal and for Dorsal/Twist cooperativity (Bhaskar *et al*. 2002; Erives and Levine 2004; Ratnaparkhi *et al*. 2008). Notch input is mediated by a Su(H) binding site as shown by over-expression of constitutively-active NICD and mutation of the Su(H) site (Markstein *et al*. 2004; Crocker *et al*. 2010). There are also conserved binding sites for the pioneer factor Zelda (Brittain *et al*. 2014), which is important for the temporal activity of many embryonic enhancers (Harrison *et al*. 2011; Nien *et al*. 2011).

Activator sites for Dorsal, Twi:Da, Dip3, and Su(H) exhibit a constrained organization in each NEE (Erives and Levine 2004). Furthermore, Dorsal gradient readouts by NEEs are sensitive to the length of a spacer element that separates the Dorsal and Twi:Da binding sites (Crocker *et al*. 2008, 2010; Crocker and Erives 2013), and is exploited in the evolutionary tuning of gradient responses (Crocker *et al*. 2008, 2010; Brittain *et al*. 2014). This functional spacer sensitivity of NEEs may involve the poly-glutamine (pQ) enriched trans-activation domains of NEE activators: Dorsal, Twi, Da, Su(H), and Zelda (see Fig. 1A). Additionally, Dorsal:Twi co-activation involves the SUMOylation system (Bhaskar *et al*. 2002; Ratnaparkhi *et al*. 2008), which can attenuate pQ-mediated aggregation (Mukherjee *et al*. 2009). The carboxamide side-chains of both glutamine (Q) and asparagine (N) participate in additional hydrogen bonding, which is a key feature of pQ/pN-mediated protein aggregation. Although proteins with pathological expansions of pQ tracts ≥ 40 residues can self-assemble into cross-*β*-sheet amyloid fibers (Perutz *et al*. 2002a,b), proteins with shorter pQ tracts can aggregate into complexes when imported into the nucleus and brought together by a DNA scaffold (Perez *et al*. 1998). As such, the lengths of functional *cis*-spacers may modulate the degree of pQ aggregation and/or *β*-strand interdigitation (Rice *et al*. 2015).

**Figure 1.**
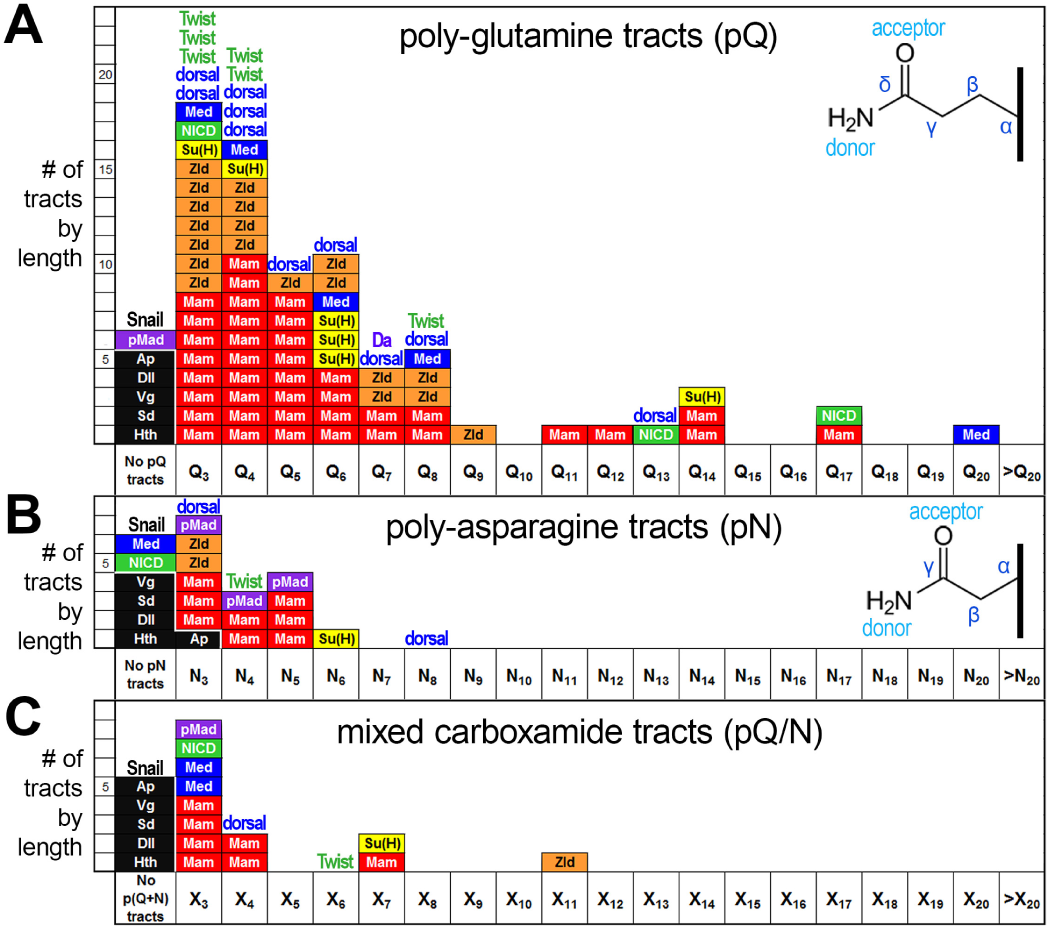
Poly-glutamine (pQ) and poly-asparagine (pN) content of factors targeting Notch-signal integrating enhancers. **(A)** Shown is a histogram of pQ tracts for the Dpp-effectors, pMad (violet) and Medea (Med, blue); the Notch-effectors, Su(H) (yellow) and its co-activators, the Notch intracellular domain (NICD, green) and Mastermind (Mam, red); and the temporal patterning factor Zelda (Zld, orange). Binding sites for these factors are enriched in the *nab* DWME. Also shown is the pQ-content for activating TFs targeting the analogous NEEs: dorsal, Twist, Daughterless (Da), Zld, and Su(H). Snail (Sna) is a NEE-targeting transcriptional repressor and is devoid of pQ content. Each pQ tract, defined as a contiguous sequence of Q’s ≥ 3, is represented by a single box in a bin corresponding to its length. In contrast, to the patterning factors, the selector DNA-binding factors (Ap, Dll, Sd, Hth) and co-factors (Vg) are devoid of pQ tracts likely indicating distinct modes of regulation separate from the signaling pathway effectors (see text). Similar trends are also seen with poly-asparagine (pN) **(B)**, or with mixed poly-carboxamide (X) tracts **(C)**.

In *D. melanogaster*, the NICD co-activator contains a single, long, nearly uninterrupted pQ tract (“Q13HQ17”) that is conformationally variable and functionally important (Wharton *et al*. 1985; Kelly *et al*. 2007; Rice *et al*. 2015). Discovery of the Notch pQ tract led to the study of many such pQ repeats in transcriptional activators, co-activators, and their role in synergistic activation (Wharton *et al*. 1985; Courey and Tjian 1988; Courey *et al*. 1989). Su(H)-bound NICD recruits the pQ-rich Mastermind co-activator (Yedvobnick *et al*. 1988; Newfeld *et al*. 1993; Helms *et al*. 1999; Schuldt and Brand 1999; Kovall 2007), as well as pQ-rich Mediator components (Janody and Treisman 2011; Tóth-Petróczy *et al*. 2008). Thus, Notch-permissive enhancers might function through pQ-aggregated complexes that accumulate in the nucleus. These enhancer-specific complexes might continue forming for the duration that NICD and other pQ-rich signals are being received by these enhancers. To understand the logic and functional repertoire of Notch-permissive enhancers, we seek to identify and study novel Notch target enhancers that also read-out morphogenic signals via pQ-rich effectors.

Here we identify and analyze a novel, Su(H)-dependent, Notch/BMP-integrating enhancer in the BMP-inducible gene *nab*, which encodes a conserved transcriptional co-factor (Clements *et al*. 2003; Terriente Félix *et al*. 2007; Ziv *et al*. 2009; Hadar *et al*. 2012). The *nab* enhancer drives expression in two domains abutting the dorsal wing margin and flanking the stripe of Dpp expression in *Drosophila*. We find that this dorsal wing margin enhancer (DWME) is licensed by the selector Apterous (Ap) to read-out Notch and BMP/Dpp signaling in the dorsal compartment of wing imaginal discs. Ap is a homeodomain containing factor specifying the dorsal compartment of wing imaginal discs (Cohen *et al*. 1992; Blair *et al*. 1994). We show that the activity of the *nab* DWME, which has multiple Ap-binding sites, is affected by the dosage of *Ap*, and find that *nab* is required in the dorsal compartment for morphogenetic patterning of the thorax and wings. We show that activity of the *nab* DWME is driven by Notch signaling through at least two Su(H) binding sites, and discuss how it likely functions as a lineage-specific enhancer in which BMP effectors maintain DWME activity in off-margin descendants of NICD-positive margin cells. Last, we show that global *nab* expression is driven by the combined activity of the *nab* DWME, a wing pouch quadrant enhancer (QE), a proximal wing enhancer (PWE), and a larval brain enhancer (BrE), all of which function as dual enhancers and silencers. Importantly, we find that some of their Su(H) sites function in inter-enhancer silencing, revealing an important hidden aspect of Su(H)-targeted enhancers.

## Materials and Methods

### Derivation of binding motifs

Derivations of IUPAC DNA motifs from position-weighted matrices from various data sets are illustrated in Fig. S1 (Supporting File S3, also see Table 1). The Mad:Medea binding motif 5′-CNBYGDCGYSNV is a consensus IUPAC motif that we derived from embryonic ChIP-chip data (Bergman *et al*. 2005) available on JASPAR (Sandelin *et al*. 2004; Mathelier *et al*. 2015). The Zelda binding motifs, 5′-CAGGYAR,5′-CAGGTAV, and 5′-YAGGTAR, were based on previously published characterization of Zelda ChIPchip data (Harrison *et al*. 2011; Nien *et al*. 2011). The Apterous binding motif 5′-RYTAATKA is a consensus IUPAC motif, which we derived from bacterial 1-hybrid (B1H) data from the FlyFactorSurvey project (Zhu *et al*. 2011). We also derived the Pangolin (dTcf) binding motif 5′-TTTGWWS from B1H data available from FlyFactorSurvey. For the Scalloped dimer binding motif, which is necessary for recruitment of Vestigial, we re-aligned the known binding sites from the *cut*, *sal*, and *kni* enhancers (Halder and Carroll 2001), and found we could derive the tighter consensus 5′-RVATTNNNNRVATH by using the reverse complement sequences of some sites in a consensus alignment (see Fig. S1 in Supporting File S3). Sequences containing the above sites were identified from the 1344 conserved regulatory belts containing sites matching the Su(H) motif 5′-YGTGRGAAH. Motifs were searched using Regular Expression pattern matching, perl, and grep commands in a UNIX shell environment.

### Molecular cloning

DNA fragments were amplified from either genomic DNA extracted from *w*^1118^ flies (DNA ⇒ *nab-C*, and DNA ⇒ *nab-A* ⇒ *nab-Ax*), or DNA prepared from the BDGP BAC clone BACR48M07 and its sub-cloned derivatives (BAC ⇒ *nab-CDAB* ⇒ *nab-AB* ⇒ *nab-B*, *nab-CDAB* ⇒ *nab-DAB* ⇒ *nab-DA*, and BAC ⇒ *nab-CDA*). All enhancer fragments were sequenced in both directions to confirm identity of clones and the absence of unwanted mutations, although the larger fragments were not necessarily sequenced through their entire lengths (see Supporting Files S1 and S2). Su(H) mutations were created using two-step PCR-mediated stitch mutagenesis to introduce changes as indicated in the text, and sequenced to confirm these mutations. Outside of the introduced mutations, the sequences obtained for mutagenized clones are otherwise identical to their parent clones (*nab-A*, *nab-AB*, or *nab-CDA*). To sequence inserts cloned into the pH-Stinger vector we used Stinger-FWD: (5′ATA CCA TTT AGC CGA TCA ATT GTG C) and Stinger-REV: (5′CTG AAC TTG TGG CCG TTT ACG).

Amplified *nab* intronic fragments were cloned into the Xba I site of the pH-Stinger vector (TATA-box containing *hsp70* core promoter driving nuclear eGFP) (Barolo *et al*. 2004). The *nab A* fragment was also cloned into the EcoR I site of the “-42 eve-lacZ” pCASPAR vector. The *nab* intronic fragments were amplified using a high-fidelity *Taq* polymerase (NEB Platinum Taq mix) and the following oligonucleotide primer pairs designed from the reference iso-1 assembly: *nab-A*: A-fwd (5′-TGGACGCAACTGGTCTGATA) and A-rev (5′-GACCAAGGATGCGATACGAT); *nab-B*: B-fwd (5′TTTCAGAAGGGGTTGAACC) and B-rev (5′-CGTATGCATAAGAAACTGGC); *nab-C*: C-fwd (5′-ACAAGTACAATGGACATGG) and C-rev (5′GAAAAGATACATATGAGTAATGC); *nab-Ax*: A-fwd and Ax-rev (5′-CCAGCAAGGATTGCCAGG); *nab-AB*: A-fwd and B-rev; *nab-DA*: D-fwd (5′-CTCATATGTATCTTTTC, which spans reverse complement of C-rev) and A-rev; *nab-DAB*: D-fwd and B-rev; *nab-CDA*: C-fwd and A-rev; and *nab-CDAB*: C-fwd and B-rev. All primers included flanking restriction sites for Xba I (5′-TCTCAGA) or Bsa I/EcoR I (5′-GGTCTCGAATTC).

### Dissection and Antibody Staining

Wandering third instar larvae were dissected and fixed with 11.1% formaldehyde in PBS for 30 minutes. Tissue was washed for 30 minutes in PBT and then blocked with 1% BSA in PBT for 1 hour. Tissue was incubated with primary antibodies over night and with secondary antibodies for 1.5 hours. After each antibody incubation, a series of washes was done for 30 minutes. Nuclear stained tissue was incubated with 17.5*µ* M DAPI in PBT for 5 minutes, and then washed for 30 minutes. Subsequently the imaginal discs or larval brains were dissected from the remaining tissue and cuticle. This dissection was done on a slide in 80% glycerol. The slide was then covered with a supported cover slip and imaged with a confocal microscope.

The following primary antibodies were used: chicken anti-GFP (1:250) (abcam: ab13070), rabbit anti-Gal4 (1:250) (Santa Cruz Biotechnology: sc-577) mouse anti-Wg (1:25), mouse anti-Ptc (1:50), mouse anti-En (1:50), and mouse anti-*β*galactosidase/40-1a (1:12). The last four monoclonal antibody were developed by Cohen, S.M., Guerrero, I., Goodman, C. and Sanes JR. (respectively) and were obtained from the Developmental Studies Hybridoma Bank as concentrates/supernatant and developed under the auspices of the NICHD and maintained by The University of Iowa, Department of Biology, Iowa City, IA 52242. The primary antibodies were detected with Cy2–conjugated Goat anti-Chicken (1:1000) (abcam:ab6960), Cy3-conjucgated Goat anti-Rabbit (1:1000) (abcam:ab6939) or Cy5-conjugated Goat anti-Mouse (1:1000) (Invitrogen:A10525) secondary antibodies.

### Drosophila Stocks

Functional analysis of *nab* and the *nab* DMWE were performed using stocks obtained from the Bloomington, Kyoto, and VDRC stock centers. The stocks from the Bloomington Stock Center are as follows: *N*^1^ (6873), *ap^Gal^*^4^ (3041), enGal4 UAS:Dcr (25753), UAS:GFP (6874). The *nab*-RNAi (1607) stock was obtained from the VDRC stock center. The following *nab* enhancer trap stocks were obtained from the Kyoto Stock Center: nab-NP1316 (112622) and nab-NP3537 (104533).

### Notch Mutant Cross

Virgin N^1^/FM7c flies were crossed with *nab DWME* (Xchromosome) males. Overnight embryos were collected and sorted by GFP (corresponding to the twist:Gal4 UAS:GFP from the FM7c balancer) at 0, 12, and 24 hours after collection. Sorted embryos grown to wandering third instar were dissected and stained as detailed above.

### Data Availability

Transgenic lines carrying all *nab* enhancer reporters described in this study are available upon request.

Supporting File S1 is a text file containing FASTA sequences for the entire *nab-CDAB* enhancer complex (reference iso-1 genome) as well as the cloned *nab-A* sequence. Annotated sequences for the *nab*-CDAB fragment and the cloned *nab*-A fragment sequences have been deposited at GenBank under the following accession numbers (TO BE ADDED WHEN RECEIVED). Supporting File S2 is a pdf file showing an annotated alignment of all cloned sequences relative to the reference genome. Supporting File S3 contains four supporting figures as follows. Figure S1 shows our derivation of IUPAC transcription factor motifs that we used in this study. Figure S2 shows images of *nab* DWME-driven GFP expression in live larvae and pupae and in live dissected imaginal discs. Figure S3 shows overnight beta-gal staining of wing discs dissected from all seven independent P-element lines that we established for the *nab-A* (DWME) *lacZ* reporters. Figure S4 shows embryonic fixed nucleosome positions for the *nab* DWME based on data from (Langley *et al*. 2014).

## Results

### Identification of conserved regulatory belts with binding sites for Notch/BMP effectors and wing disc selectors

The Su(H) binding motif (5′-YGTGRGAAH) is remarkably constant across bilaterians and unique to Su(H) (Tun *et al*. 1994). To identify novel enhancers integrating Notch signaling with pQ-rich morphogenic effectors akin to the neurogenic ectoderm enhancers (NEEs), we first developed a computational pipeline that identified 1344 evolutionary-conserved regulatory belts containing one or more adjacent Su(H)-binding sites (pipeline and other identified enhancers will be described in a separate manuscript). We find that most regulatory belts range from 15 to 25 peaks of genus-wide conservation. Each peak typically corresponds to one to three overlapping binding sites for transcription factors (TFs). This is consistent with enhancers requiring multiple *cis*-elements for both nuanced activity patterns and restricted tissue-specificity. Here, we demonstrate how these candidate Su(H)-targeted genetic elements can be used to find model enhancers that are well suited for testing hypotheses about functional enhancer grammar.

To find enhancers that are integrating both Notch and Dpp/BMP signaling, we searched for the subset of 1344 *D. melanogaster* regulatory belts that also contain a binding site for the pQ/pN-rich Mad:Medea complex, which mediates activation of Dpp/BMP targets (Sekelsky *et al*. 1995; Newfeld *et al*. 1996; Wiersdorff *et al*. 1996; Newfeld *et al*. 1997; Hudson *et al*. 1998; Wisotzkey *et al*. 1998; Campbell and Tomlinson 1999; O’Connor *et al*. 2006; Weiss *et al*. 2010). Specifically, we developed and searched for the subset of belts containing the IUPAC consensus motif that we derived for phosphorylated-Mad (p-Mad) ChIPseq peaks (see Material and Methods and Table 1). We also searched for sequences targeted by Zelda, which is a pQ-rich pioneer factor for embryonic enhancers that is also expressed in wing imaginal discs (Staudt *et al*. 2006; Harrison *et al*. 2011). Altogether, we found 98 unique regulatory belts containing binding sites for this pQ-rich set of activators Su(H), p-Mad:Medea, and Zelda (Table 1 and Fig. 1).

To find enhancers that are specific to the wing imaginal disc, we refined the subset of 98 down to those that also had binding sites for three wing imaginal disc selectors: **(*i*)** Apterous (Ap), which is expressed throughout the dorsal wing compartment (Cohen *et al*. 1992); and **(*ii*)** Scalloped (Sd) and Vestigial (Vg), which are expressed at the dorsal/ventral compartment boundary where there is active Notch-Delta signaling (Halder *et al*. 1998; Halder and Carroll 2001; Koelzer and Klein 2006). By definition, selectors are TFs that are responsible for a stable binary cell fate decision (García-Bellido 1975; Lawrence *et al*. 1979; Akam 1998). For Ap, we derived the IUPAC consensus motif 5′-VYTAATKA from its DNA binding profile and found 31/98 regulatory belts with sequences matching this motif (Table 1). For Sd and Vg, we found that 13 of these 31 regulatory belts contain the canonical Sd dimer binding site that recruits the Sd:Sd:Vg complex (Table 1) (Halder and Carroll 2001). All three selectors (Ap, Sd, and Vg) are deficient in pQ/pN tracts unlike the graded signal-dependent effectors (*e.g*., see Fig. 1), Mediator co-activators subunits, and TBP.

Finally, to focus on pQ-rich graded signal integration of just the Notch and BMP pathways, we set aside 11 of the 13 regulatory belts that had potential binding sites for Tcf/Pangolin. Tcf/Pangolin mediates transcriptional activation in the Wnt and *β*-catenin/Armadillo signaling pathway (Van de Wetering *et al*. 1997; Archbold *et al*. 2014). This last filtering step left us with two regulatory belts at the *nab* and *crossveinless c* (*cv-c*) loci, which are located on the left and right arms of chromosome three, respectively. The gene *nab* is a BMP-induced transcriptional co-factor involved in embryonic and imaginal disc development (Clements *et al*. 2003; Terriente Félix *et al*. 2007; Ziv *et al*. 2009; Hadar *et al*. 2012), while the gene cv-*c* encodes a BMP-induced Rho-type GTPase activating protein (RhoGAP) involved in cross-vein morphogenesis (Matsuda *et al*. 2013). We later describe how RNAi knock-down of *nab* phenocopies *crossveinless*-type wings, suggesting both *nab* and *cv-c* function in a common developmental genetic pathway. In this study, we focus on the *nab* regulatory belt and its interactions with adjacent regulatory elements because it has more sites for each factor than the *cv-c* regulatory belt (Table 1).

### The nab dorsal wing margin enhancer (DWME) drives D/V wing margin expression modulated along the A–P axis

The dense site cluster overlying the *nab* regulatory belt is located in the long first intron of *nab* in *Drosophila* (thick bar underline in Fig. 2A). This same belt is part of a smaller 2.7 kb intronic fragment containing several regulatory belts conserved in both *D. melanogaster* and *D. virilis* (see boxed region in Fig. 2A). We cloned the predicted 764 bp “*nab-A*” fragment and found it corresponds to a dorsal wing margin enhancer (DWME) with augmented expression in regions corresponding to peak p-Mad levels (Fig. 2B–L). (Note: this fragment is 764 bp long in the reference iso-1 genome, but is 762 bp long in the cloned fragment due to polymorphic indels in two separate poly-A tracts.) The *nab-A* fragment was tested using two different core promoters with different reporter transgenes: **(*i*)** a 270 bp minimal *hsp70* core promoter-eGFP-nls fusion, and **(*ii*)** a 205 bp minimal *eve* core promoter-*lacZ* fusion (Fig. 2C). These constructs also tested the *nab-A* enhancer fragment in both orientations relative to these core promoters (Fig. 2B,C). We generated seven and five independent P-element integrations of the lacZ and eGFP-nls reporter constructs, respectively, and found all twelve to drive the same expression pattern.

**Figure 2.**
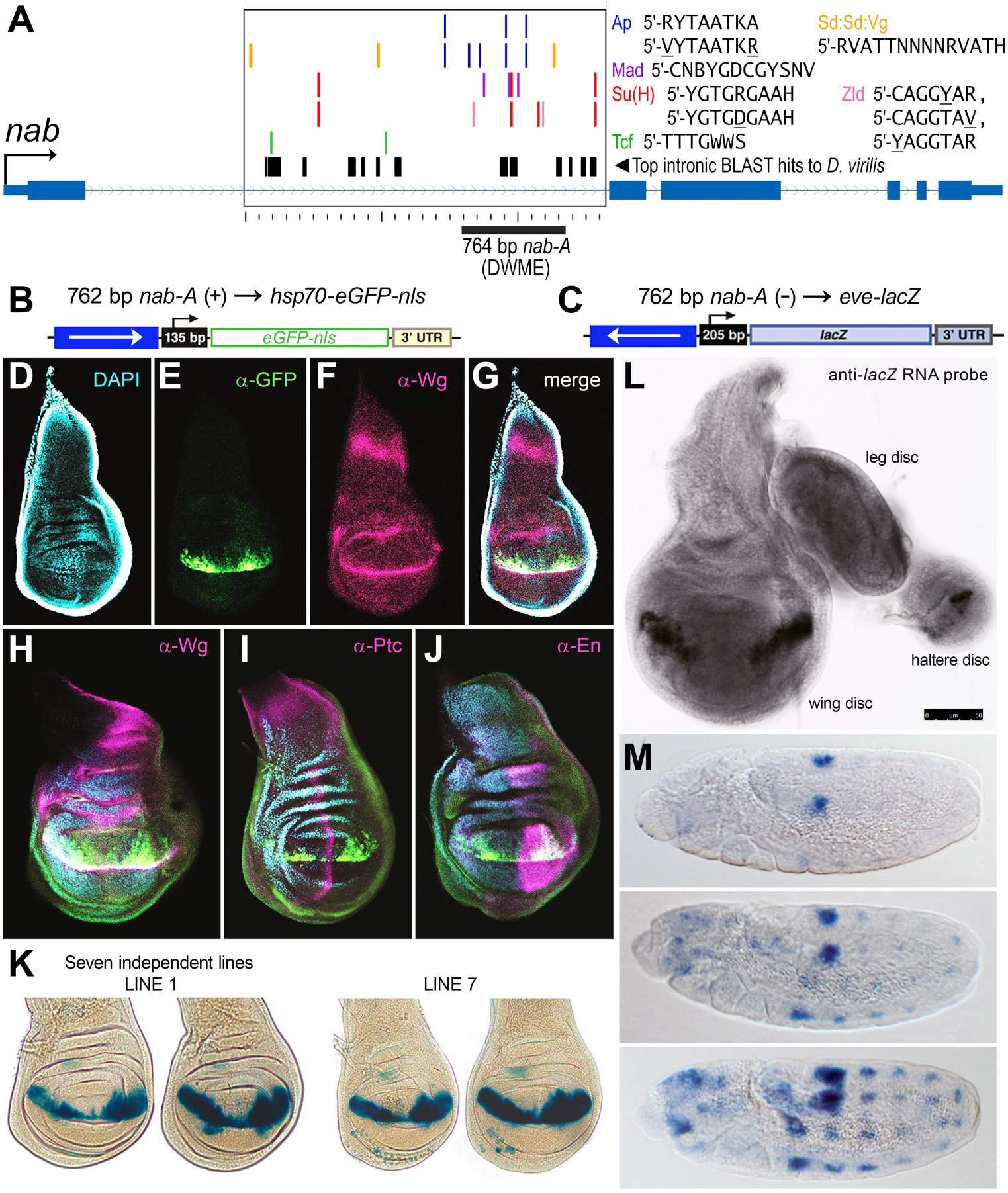
A highly-conserved regulatory block in the *Drosophila nab* locus is a robust dorsal wing margin enhancer (DWME). **(A)** Diagram of the *nab* locus from *D. melanogaster* showing sites matching binding motifs (color-coded boxes on separate tracks in large intronic box) within a 2.7 kb intronic block that contains several regulatory belts conserved across the genus. We cloned a predicted 764 bp region, the *nab-A* fragment (thick bold underline), for having one of the highest concentration of sites for Su(H), Mad, Zelda, Apterous (Ap), and the Scalloped (Sd) and Vestigial (Vg) complex (Sd:Sd:Vg) without having any Tcf sites. **(B, C)** We cloned the *nab-A* fragment in front of two core promoter reporter genes: an *hsp70* core promoter fused to an eGFP-nls reporter gene in a gypsy insulated construct **(B)**, and an *eve* core promoter fused to the *lacZ* reporter gene **(C)**. Independent P-element transgenic lines with each reporter cassette gave identical expression patterns despite differences in integration sites, core promoters, and enhancer orientation relative to the core promoter (arrows in enhancer box, and + or − signs in parentheses). The cloned 762 bp fragment is 2 bp shorter than the reference sequence due to single bp contraction polymorphisms in two separate poly-A runs. **(D–F)** Panels show the expression from the *nab-A* (DWME) EGFP reporter along the dorsal wing margin. Different color channels indicate DAPI (D, cyan); GFP (E, green); and Wingless (Wg) (F, magenta), which marks the D/V compartment margin, a ring around the wing pouch, and a broad stripe across the proximal part of the wing disc. **G** Panel shows merged imaged of D–F. **(H–J)** Additional discs are shown double-labeled with antibodies to Wg (H), and Ptc (I) and En (J), which mark the A/P margin and posterior compartment, respectively. The anterior DWME expression pattern is characteristically longer than the posterior compartment side, which in turn stretches deeper into the dorsal compartment than the anterior compartment expression pattern. **(K–M)** Panels show DWME-driven *lacZ* reporter activity in late third instar discs **(K, L)** or in germ-band extended embryos **(M)**. DWME-driven expression was detected with over-night X-gal staining (K) or with a digU-labeled anti-sense *lacZ* RNA probe (L, M). Both wing (K, L) and haltere (L) discs show the characteristic twin spots of DWME-driven activity, while the embryonic expression is detected in a subset of lateral neuroblasts.

We find that the *nab-A* fragment works equally well across these different reporter constructs in live dissected and undissected discs (Fig. S2 in Supporting File S3), in fixed and double-stained imaginal discs (Fig. 2D–J), in fixed discs stained for *β*-gal expression (Fig. 2K and Fig. S3 in Supporting File S3), and in fixed discs incubated with an anti-sense *lacZ* RNA probe (Fig. 2L). In situ hybridization with the same anti-sense *lacZ* RNA probe also shows that the *nab-A* fragment drives expression in stage 9/10 long-germ band extended embryos in what may be a subset of embryonic neuroblasts, a known site of *nab* expression (Clements *et al*. 2003).

In the wing imaginal disc, the *nab* DWME drives expression from the D/V border region and into the dorsal compartment by several cells but only in two spots centered in the anterior and posterior compartments, which correspond to regions of peak p-Mad levels. The *nab* DWME also drives expression in a highly stereotyped pattern that is unique to each compartment. In addition to the thin interrupted row of expression along the D/V margin, expression occurs as an elongated anterior spot and a broader mitten-shaped posterior spot abutting the margin. This robust DWME-driven expression pattern is consistent with our desired goal of finding a wing disc compartment-specific enhancer integrating two orthogonal developmental signals: **(*i*)** a dorsal/ventral (D/V) Notch margin signal, and **(*ii*)** a graded anterior to posterior (A–P) BMP morphogenic signal. Because endogenous *nab* is expressed in a broader wing pouch pattern than what we observe for the *DWME*-driven reporters, below we address how the DWME relates to other possible enhancers at *nab*. We also address how some of the predicted sites contribute to DWME activity, and how the DWME contributes to Nab function in wing development.

### The first intron of nab harbors four separate enhancers

To understand the role of the *nab* DWME activity in the overall *nab* expression pattern, we made a series a constructs containing various intronic fragments that we cloned from the first intron of the *nab* locus (Fig. 3A). These were tested by anti-GFP antibody staining of dissected larval tissues from independent P-element integrated lines as well as pools of these lines (see Materials and Methods). Consistent with the presence of insulators in the pH-Stinger vector, independent lines of the same construct recapitulated each other’s expression patterns.

**Figure 3.**
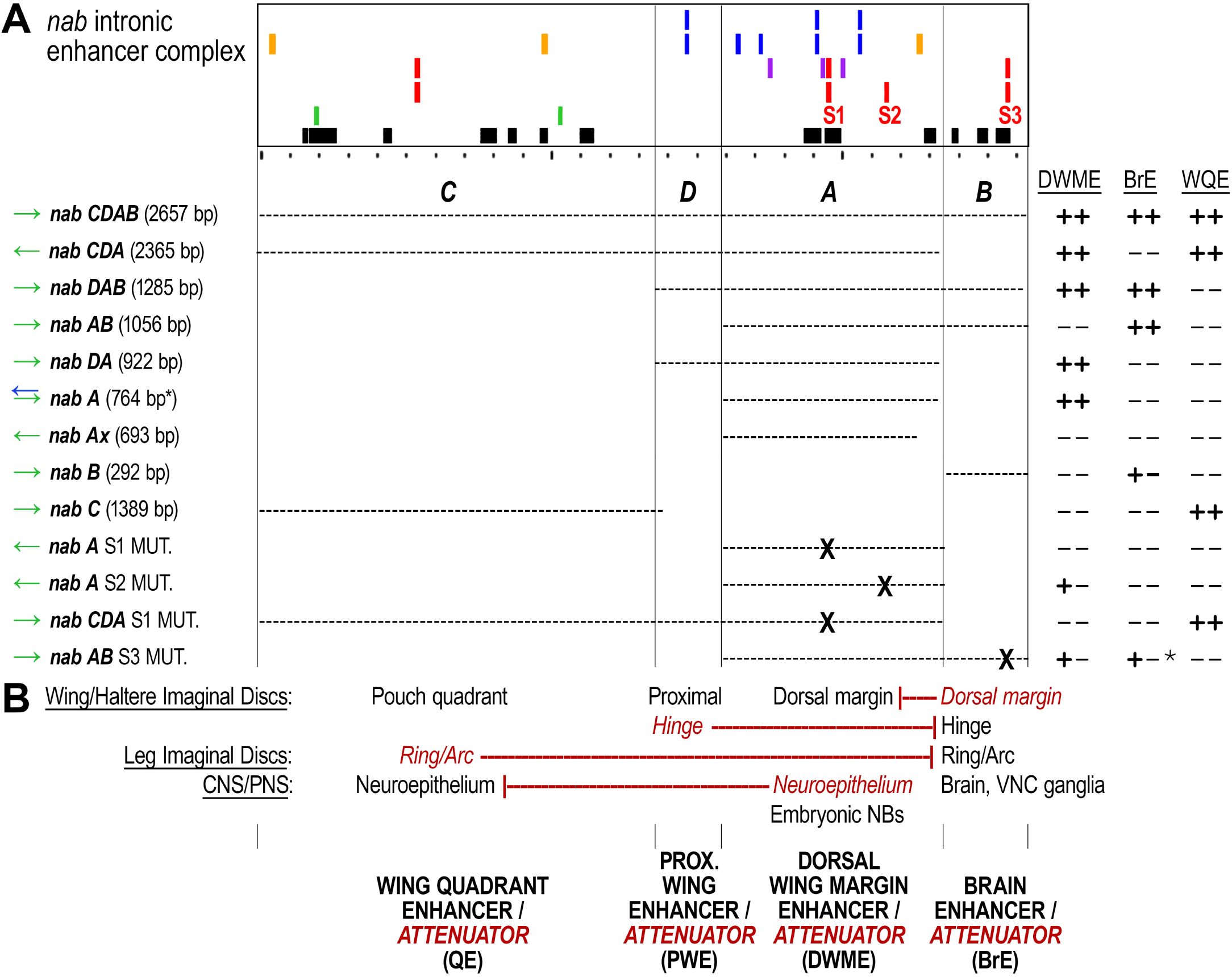
The *Drosophila nab* harbors several regulatory modules functioning as both enhancers and mutual silencers/attenuators. **(A)** We cloned and tested the indicated series of *nab* intronic fragments in order to understand the regulatory logic of *nab*’s expression in wing imaginal discs. The colored boxes follow the key in Figure 2 except Zelda sites are not shown for clarity. To refer to the different enhancers we identified, we defined four different intronic regions as the *nab-C*, *nab-D*, *nab-A* (DWME), and *nab-B* fragments. We also labeled the four best matches to Su(H) binding motifs (sites S1–S3) and mutated these sites in the indicated constructs (S# MUT., and X’s in construct). These regions can be understood as having four major enhancer activities: the dorsal wing margin enhancer (DWME); a larval brain enhancer (BrE), which drives dense expression in neuronal lineages, a wing imaginal disc quadrant enhancer (QE), which complements the DWME activity to match the endogenous expression pattern; and a proximal wing enhancer (PWE). Two plus signs indicates strong expression, one plus sign indicates weak expression while maintaining the indicated pattern. An asterisk means there is a more nuanced description of the expression in the main text. Both the sizes and the direction of the cloned insert relative to the core promoter are shown for each construct. **(B)** Each cloned enhancer was found to drive a distinct expression activity associated with the endogenous *nab* locus, a distinct ectopic activity or expression level that was not associated with the endogenous *nab* locus, and a distinct silencing or attenuation activity acting on ectopic expression patterns and levels (red repression symbols connecting one silencer/attenuation activity in one enhancer to the ectopic activity/level in another intronic enhancer).

After comparing a PGawB *nab GAL4* enhancer trap line (*nab^NP^*^1316^, insertion site 48 bp upstream, Hayashi *et al*. (2002)) with the various constructed transgenic reporter lines, we discovered that we identified separate enhancer activities for all known tissues in which *nab* is expressed (Fig. 4A panels versus B–I panels). The expression pattern of the endogenous *nab* locus in late third instar wing imaginal discs (Fig. 4A) is driven by a combination of: **(*i*)** the DWME activity from the 764 bp *nab-A* fragment (Fig. 4B), **(*ii*)** a quadrant enhancer (QE) activity from a 1.4 kb *nab-C* fragment (Fig. 4C), and **(*iii*)** a proximal wing disc enhancer (PWE) activity from the combined 2.4 kb *nab-CDA* fragment (triple arrows pointing to dispersed expression pattern in Fig. 4D). The PWE activity, which we also find in both the endogenous enhancer trap line (see triple arrows in Fig. 4A) and a larger 2.7 kb *nab-CDAB* fragment (Fig. 4E), likely requires elements present in both *nab-C* and *nab-D* fragments because neither the *nab-C* (Fig. 4C) nor the *nab-DA* fragment (Fig. 4H) recapitulates this pattern on its own.

**Figure 4.**
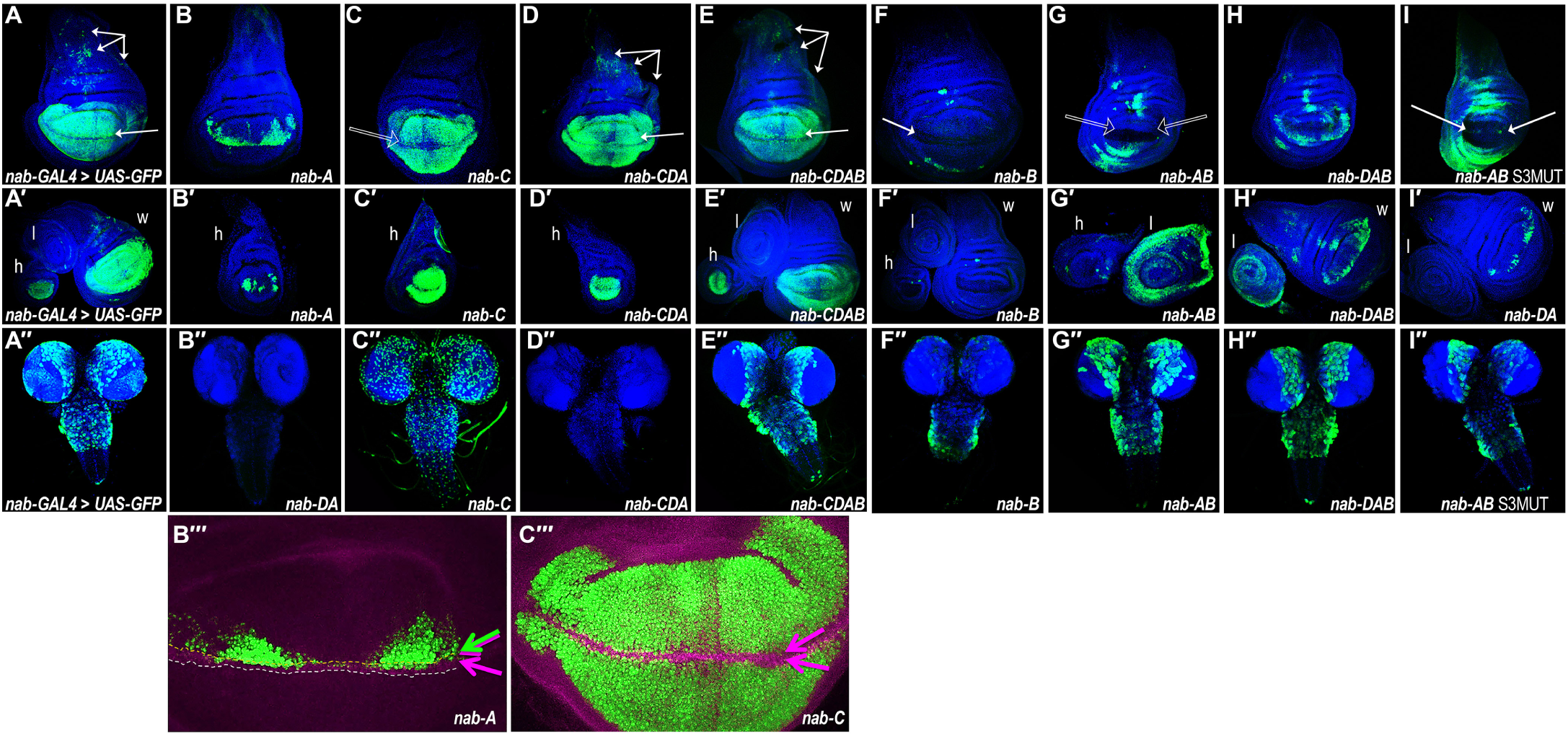
Activities of dissected regulatory modules from the *nab* intronic enhancer/silencer complex. **(A–I)** GFP reporter expression driven by *nab-GAL4*>*UAS-eGFP* **(A)** or by different *nab* enhancer modules driving *eGFP-nls* **(B–I)**. The first row depicts third instar wing imaginal discs, while the middle row (single prime) depicts additional third instar discs labeled for halteres (h), leg (l), and wing (w) discs. The third row (double prime) depicts expression patterns in third instar larval brains, if any. **(A–I)** The endogenous *nab* expression pattern in wing imaginal discs **(A)** appears to be the result of discrete activities from: the *nab-A* fragment, which contains the dorsal wing margin enhancer **(B)**; the *nab-C* fragment, which contains a wing pouch/quadrant enhancer **(C)**; a nondescript proximal wing disc enhancer seen with most intronic fragments containing the *nab-D* region (triple arrows in D, E); and the *nab-B* fragment, which contains a hinge enhancer that functions in the absence of the *nab-C* fragment (F, G, H, I). Note that the *nab-C* fragment is missing expression at the margin and has more uniform levels of expression throughout the pouch relative to either the *nab-GAL4* enhancer trap **(A)** or fragments containing both the *nab-C* and *nab-A* regions (empty arrow in C). **(A**′**–I**′) Fragments with activities from both the *nab-A* and *nab-C* fragments recapitulate in endogenous *nab* expression in the haltere disc. **(A**″**–I**″) Most of the endogenous expression of *nab* in the larval brain and ventral nerve cord **(A**″) is recapitulated by *nab* intronic fragments carrying the *nab-B* fragment **(E**″**–H**″) but it is noticeably weaker by itself **(F**″). This pattern corresponds to dense expression in neuronal lineages and four cells in the posterior tip of the ventral nerve cord. The *nab-C* fragment has ectopic brain activity that is completely silenced by activities present in the *nab-DA* fragment (see entire brain in D′ and optic lobes and posterior ventral nerve cord regions in E″). The *nab-AB* fragment also drives a strong ectopic leg disc ring pattern (see G′ and H′) that is repressed in the presence of the *nab-C* fragment (see E′). Similarly, the dorsal wing margin activity of *nab-A* **(B)** is silenced in the presence of *nab-B* (see empty arrows pointing to margin in *nab-AB* disc in G) except when *nab-D* is also present (see *nab-DAB* disc in H). The S3 Su(H) binding site in *nab-B* appears to mediate repression of the wing hinge activity inherent to *nab-B* and the dorsal wing margin activity inherent to *nab-A* because there is augmentation of both of these patterns in the *nab-AB* S3 mutated construct (I, arrows point to twin spots along the dorsal margin of the compartment boundary). This same site is not absolutely required for the *nab-B* brain activity **(I**″). Neither the *nab-DA* nor the *nab-A* are able to drive any brain expression (representative blank *nab-DA* disc is shown in B″), strongly suggesting that the *nab-B* fragment is necessary and sufficient for the overall gross expression pattern of *nab* in the brain. (**B**″′**,C**″′) Shown are blown-up images of the *nab-A* and *nab-C* activities in the wing disc (green) double labeled for Wg (magenta).

The robust expression of the endogenous enhancer trap reporter in the dorsal margin of the wing pouch (single horizontal arrow in Fig. 4A) is recapitulated in all reporter lines carrying the *nab-A* fragment (Fig. 4B and single horizontal arrows in Fig. 4D,E) with one exception summarized in the next section. Furthermore, this dorsal margin activity is absent in the *nab-C* fragment, which only drives a quadrant pattern in wing imaginal discs (see hollow arrow pointing to gap of expression at margin in Fig. 4C). The combined *nab* QE+DWME expression pattern, which we see in the *nab-CDA* and *nab-CDAB* reporters and the *nab^NP^*^1316^ P{GawB} enhancer trap (Fig. 4A, D, and E), is also recapitulated by a second P{GawB} enhancer trap (*nab^NP^*^3537^, data not shown), which has an insertion site one base pair upstream of *nab* but in the opposite orientation as *nab^NP^*^1316^, and a third, previously characterized P{GabB} *nab* enhancer trap line, S149, which has a reported insertion site 23 bp upstream (Gerlitz *et al*. 2002; Ziv *et al*. 2009; Hadar *et al*. 2012).

We also observed that all *nab* enhancer fragments driving wing pouch expression patterns (DWME and QE) also did so in the haltere imaginal discs (“h” discs in single prime lettered panels of Fig. 4). This suggests to us that *nab* plays a role in both wing and haltere development that predates evolution of haltere balancing organs from wing discs in dipteran ancestors (Lewis 1978; Weatherbee *et al*. 1999).

The endogenous enhancer trap reporter is expressed in entire neuronal lineages of the larval brain and in four distinct neurons/cells at the posterior tip of the ventral nerve cord (Fig. 4A″). We find that a 292 bp *nab-B* fragment, located immediately downstream of *nab-A* (Fig. 3A), drives the same pattern but at lower levels than the endogenous locus (Fig. 4F″). These lower levels of expression are rescued by larger fragments containing the *nab-B* region: *nab-AB* (Fig. 4G″), *nab-DAB* (Fig. 4H″), or *nab-CDAB* (Fig. 4E″) fragments. This suggests that the minimalized larval brain enhancer (BrE) present in the *nab-B* fragment normally functions with additional elements that span into the adjacent *nab-A* fragment. Nonetheless, a 693 bp truncated *nab-A* fragment (see *nab-Ax* fragment in Fig. 3A), which is missing a conserved sequence that is present on the side flanking the *nab-B* fragment, does not drive any detectable activity in either imaginal discs or larval brain (data not shown). This might signify that the conserved block present in *nab-A* and separating it from the *nab-B* fragment is required for both the DWME and BrE activities. Alternatively, BrE boosting elements may be located elsewhere in the *nab-A* fragment.

**Table 1.**
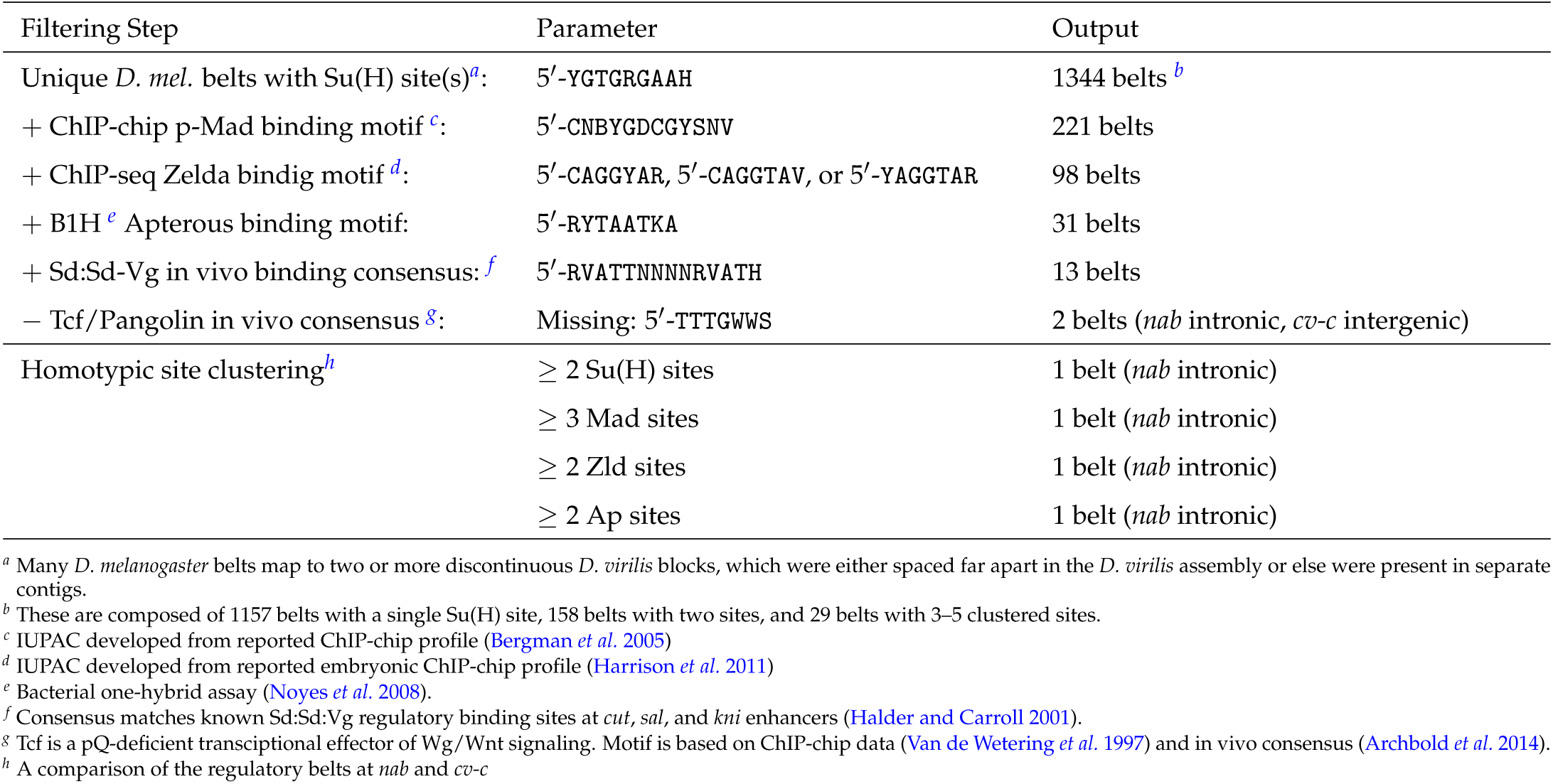
Regulatory belts with sites for pQ-rich effectors of Notch/BMP signaling and pQ-deficient wing disc selectors.

### The nab enhancers also function as mutual silencers

For each of the four distinct and separable enhancer activities in the first intron of *nab*, we also find distinct silencer activities (Fig. 3B). For example, while the *nab-C* fragment does not recapitulate the larval brain enhancer (BrE) activity, it does drive ectopic expression in a more superficial set of larval brain cells, possibly neuroepithelial cells or glia, and some distinct neurons in the ventral nerve cord (Fig. 4C″). This ectopic activity associated with *nab-C* is not seen from the endogenous locus (Fig. 4A″) or from any fragment that also contains the *nab-DA* region (Fig. 4D″,E″). This suggests that the ectopic larval brain activity endowed by *nab-C* is silenced by elements present in *nab-DA* (Fig. 3B).

In a second example, we find that the *nab-AB* enhancer is a potent ring enhancer in leg imaginal discs (“l” disc in Fig. 4G′), which is an activity that is not seen for the *nab* enhancer trap reporter (“l” disc in Fig. 4A′). This activity is attenuated (*i.e*., made less robust) in the *nab-DAB* fragment (“l” disc in Fig. 4H′), and is completely silenced in the larger *nab-CDAB* fragment. This suggests that the ectopic leg disc activity endowed by *nab-AB* is silenced by elements spanning *nab-C* and *nab-D* (Fig. 3B).

In yet a third example, which we previously mentioned as an exception to the presence of DWME activity in all fragments containing *nab-A*, we find that the *nab-AB* fragment obliterates dorsal wing margin expression (arrows Fig. 4G). However, this same silencing activity is only able to attenuate the DWME in the context of the larger *nab-DAB* fragment (Fig. 4H and “w” disc in H′). These results suggest that the *nab-B* fragment is simultaneously a larval brain enhancer and a dorsal wing margin silencer/attenuator. In a related fourth example, the *nab-D* fragment also appears to modulate and attenuate *nab-A*’s DWME activity in the context of the *nab-DA* fragment (Fig. 4I).

A fifth example of dual enhancer/silencer activity is intertwined with the above third example of the BrE’s wing disc silencing activity. The *nab-AB* fragment drives ectopic expression in the wing imaginal disc hinge region near the stripe of peak Dpp (Fig. 4G). This activity is attenuated in the *nab-DAB* fragment (Fig. 4H and “w” disc in H′) and completely silenced in the *nab-CDAB* fragment (Fig. 4G).

Thus, we find that all of the distinct *nab* enhancer fragments possess both endogenous (*i.e*., *nab*-related) enhancer activities and ectopic enhancer activities. However, these same enhancers also function as inter-enhancer silencers of ectoptic activities and attenuators that modulate their non-ectopic activities. These activities are summarized in Figure 3B. Later, we describe finding that at least one of the intronic Su(H) binding sites involved in inducible activation is also involved in inter-enhancer silencing and attenuation.

### Activity of nab DWME requires Notch signaling and Su(H) sites

Single nucleotide transversions to any of the core positions in the invariant Su(H) binding sequence, 5′-Y<u>GTG</u>RGAA (core positions underlined), are sufficient to eliminate binding in vitro (Tun *et al*. 1994). Therefore, to verify that the *nab* DWME is targeted by Notch signaling via its canonical Su(H) binding sites, we compared DWME activity from *nab-A* fragments containing either wild type Su(H) sites, a mutated S1 site, or a mutated S2 site (Fig. 5A–E).

**Figure 5.**
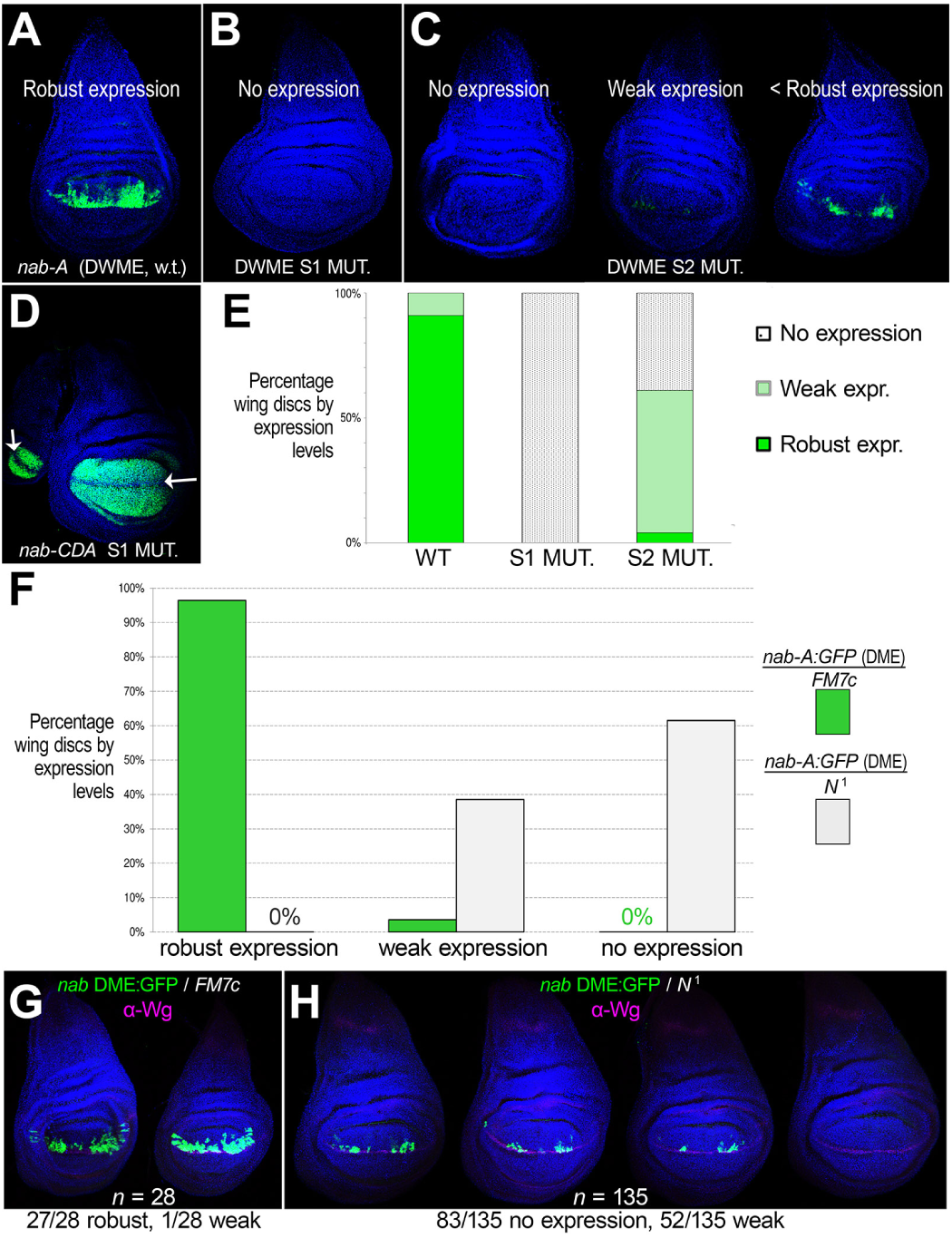
The *nab* DWME is induced by Notch signaling via its Su(H) sites. **(A)** The dorsal wing margin enhancer (DWME) contains two Su(H) sites, S1 and S2, and drives a unique stereotypical pattern. To test the role of these sites, we mutated each individually in the minimalized *nab-A* enhancer fragment. **(B)** The S1 mutated *nab-A* fragment exhibits no expression in wing imaginal discs. **(C)** The S2 mutated *nab-A* fragment results in discs with either no expression or weak dorsal margin expression as shown in these representative discs. **(D)** The *nab-CDA* fragment with an S1 mutated Su(H) exhibits reduced expression at the D/V compartment margin (arrow). **(E)** Quantitative comparison of wild type, S1 mutated, and S2 mutated discs according to levels of expression. **(F)** Quantitative comparison of *nab-A* DWME activity in wild type or mutant Notch backgrounds. Also shown is representative DWME reporter activity from discs in wild type **(G)** or mutant **(H)** Notch backgrounds.

The central S1 site is the only sequence within the *nab-A* fragment that matches the motif 5′-YGTGRGAAH, which we used in our computational screen. We found that the reporter lines with the mutated S1 site (5′-T<u>GTG</u>AGAAT → 5′-T<u>cat</u>AGAAT) lack expression in both wing and haltere discs (Fig. 5B,E). When we mutated the S1 site in the context of the *nab-CDA* fragment (Fig. 3A), we found diminished expression at the margin similar to the *nab-C* fragment (compare Fig. 5D to Fig. 4C). Thus, the Su(H) S1 site is absolutely necessary for *nab* DWME activity in both the cells along the dorsal margin and the anterior/posterior compartment cells located farther into the wing pouch.

The flanking Su(H) S2 site matches the consensus motif 5′-YGTG<u>D</u>GAAH, which is slightly less stringent than the genome screen motif 5′-YGTG<u>R</u>GAAH (Fig. 3A). Nonetheless, this S2 site is still conserved across *Drosophila*. We found that reporter lines with the mutated S2 site (5′-C<u>GTG</u>TGAAA → 5′-C<u>cat</u>TGAAA) drive diminished expression that is collapsed to two small attenuated spots along the margin near the predicted peak p-Mad regions *nab-A* (Fig. 5C,E). Thus, the flanking S2 Su(H) site is also important for robust activation along the dorsal wing margin as well as in the off-margin spots.

To confirm that the Su(H) sites work canonically in Notch-dependent inducible activation, we also compared DWME activity from the *nab-A*-driven reporter in both wild type and mutant Notch backgrounds (Fig. 5F–H). These experiments show that whereas wild type Notch allows robust DWME-activity in nearly 100% of wing imaginal discs, a Notch mutant background (*N*^1^) results in about 60% of the discs not having any expression at all and about 40% of the discs having very weak expression close to the margin (Fig. 5F, H). Thus, the *nab* dorsal wing margin enhancer (DWME) requires both wild type Notch signaling and both of its Su(H) binding sites for robust expression in the wing pouch. This signaling is also required for the off-margin expression in the anterior/posterior compartment “spots” suggesting that BMP input into the DWME is not independent of Notch signaling. These results also suggest that the Su(H) binding sites are not involved in repression in larval stages because mutating them does not uncover ectopic activation.

### Su(H) binding sites are involved in inter-enhancer silencing

In the absence of NICD, Su(H) recruits Hairless and establishes a repression complex at many targeted enhancers (Bang *et al*. 1995; Barolo *et al*. 2002; Maier *et al*. 2011). Moreover, there is evidence for long-range looping of multiple Notch/Su(H)-targeted promoters from the *E(spl)* gene complex into an interaction domain associated with cohesin and the PRC1 polycomb repression complex (Schaaf *et al*. 2013). We thus hypothesize that some of the predicted Su(H) binding sites involved in Notch-dependent inducible activation might also be involved in inter-enhancer silencing or attenuation.

To test the hypothesis that the S1 Su(H) site within the DWME is responsible for silencing the ectopic larval brain activity associated with *nab-C*, we mutated this site in the *nab-CDA* fragment (Fig. 3A). We find that mutation of this site does not result in ectopic expression in larval brains (data not shown). Nonetheless, it is still possible that the S2 site and/or additional lower-affinity sites, which we did not mutate, might participate singly or collectively in an intronic inter-enhancer repression complex featuring Su(H).

To test the hypothesis that the strong S3 Su(H) site present within the BrE is responsible for the silencing of the DWME, we mutated this site (5′-T<u>GTG</u>AGAAC → 5′-T<u>cat</u>AGAAC) in the *nab-AB* fragment (Fig. 3). Normally, the DWME activity is completely silenced in the wild type *nab-AB* fragment, and partially attenuated in the *nab-DAB* fragment. We find that mutation of this S3 Su(H) site in *nab-AB* rescues the twin spots of DWME activity in about half of the discs (16/30 discs), although at levels much diminished relative to the wild-type *nab-A* fragment (Fig. arrows 4I). Moreover, we also find that the hinge activity is much expanded along both the A–P axis and along the proximal/distal axis of the imaginal discs (Fig. 4I). This suggests that the S3 Su(H) site is involved in both intra-enhancer repression and inter-enhancer silencing/attenuation.

### Apterous licenses the nab DWME to receive Notch/BMP signals in dorsal wing compartment

We hypothesize that there are distinct mechanisms underlying binary selector “licensing” and graded signal readouts. More specifically, we hypothesize that selectors would work through nucleosomal-positioning (see Discussion), while signal pathway effectors would work through pQ-mediated aggregation and Mediator recruitment. To remain agnostic about whether instead activating TFs work through the same mechanism, we will refer to selector activity as (homeotic) licensing and the pathway effector activity as (true) activation for two reasons. First, licensing via the fixation of alternate nucleosome positions could function phenomenologically both in transcriptional activation and in repression depending on whether effector sites are being revealed or occluded with the positional shifting of a fixed nucleosome. Second, signaling pathway-based effectors tend to be pQ-rich like the Mediator co-activator complex (Tóth-Petróczy *et al*. 2008) and TATA Binding Protein (TBP) (Koide *et al*. 1999), which are recruited by pathway-effector TFs. Thus, signaling pathway integration via pQ-mediated aggregation would be inherent to eukaryotic activation and functionally-rich.

Based on the above perspective, we used the binding motifs for the pQ-deficient selectors Apterous (Ap) and others to identify the *nab* DWME. Expression of the selector Apterous (Ap) is restricted to the dorsal compartment of the wing pouch and the proximal parts of the wing imaginal disc (Cohen *et al*. 1992; Blair *et al*. 1994) (Fig. 6A). We thus hypothesize that Ap shifts or remodels a default positioned nucleosome that hides the Su(H) site in most tissues and find initial indirect support for this hypothesis in two genomic data sets for the early embryo. First, two different nucleosomal data sets for embryonic stages confirm that there are fixed nucleosomes over the DWME that we predict would obscure its Su(H) sites (Fig. S4 in Supporting File S1 shows positions found for stage 5 embryos) (Mavrich *et al*. 2008; Langley *et al*. 2014). Second, embryonic Su(H) ChIP-seq experiments (Nègre *et al*. 2011) indicate that Su(H) does not bind to the DWME during early embryonic stages, even though it is expressed ubiquitously (Schweisguth and Posakony 1992). Such a model is consistent with both recent support for a more dynamic pattern of Su(H) association with target enhancers (Krejˇcí and Bray 2007) and our nucleosome-based homeotic licensing model. We thus suggest that Ap and other selectors including Sd and Vg, may shift or remodel one or two positioned nucleosomes so as to reveal its Su(H) binding sites and permit Notch inducible activation in the dorsal wing compartment. In this regard, it is notable that Ap binding sequences coincide with nucleosomal entry sequences, which correspond to canonical linker histone H1 binding regions (Syed *et al*. 2010). In the absence of these selectors, the DWME would be refractive to Notch and possibly Dpp signaling, which are active in many contexts.

**Figure 6.**
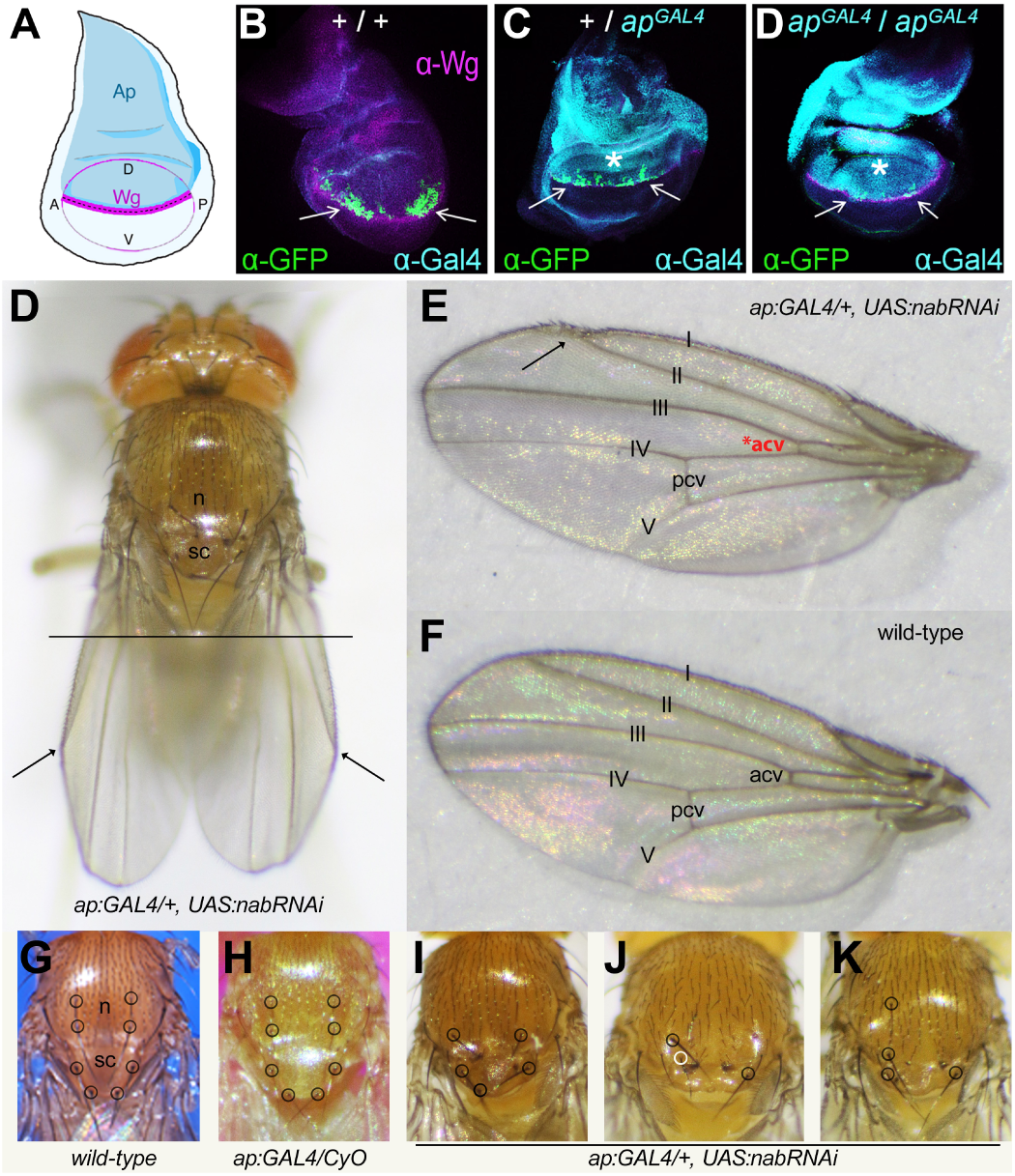
Apterous licenses the *nab* DWME to be receptive to Notch and Dpp signaling in the dorsal compartments of wing imaginal discs. **(A)** Cartoon of Ap expression (blue) in the dorsal compartment of a third instar wing imaginal disc. Wingless (Wg) expression (magenta) is found at the D/V compartment boundary in the wing pouch. **(B–D)** Wing imaginal discs stained for Wg (magenta), Gal4 (cyan), and GFP (green) in a line carrying the *nab-A* (DWME) eGFP reporter in a wild-type Ap background **(B)**, in a heterozygous *ap-GAL4* hypomorph caused by a *GAL4* element integration **(C)**, and a homozygous *ap-GAL4* hypomorph **(D)**. Note as the intensity of the Gal4 signal increases the *nab* DWME-driven GFP signal decreases, which is consistent with its many Ap binding sites. **(D,E)** The same *ap-GAL4* hypomorph was used to drive *nab* RNAi in order to knockdown expression in the dorsal compartment via both RNA and enhancer interference. Defects include an overly creased wing at wing vein II (arrows in D), a delta-like patterning-defect where wing veins I and II intersect (arrow in E), and occasionally missing or diminished anterior cross-veins (*acv). **(F)** A wild type wing for comparison to **(E)**. **(G)** A wild type notum (n) and scutellum (sc) with macrochaetes circled. **(H)** One balanced copy of the *ap-GAL4* does not affect macrochaete patterning nor development of the scuttelum. **(I– K)** When *ap-GAL4* drives *nab*-RNAi, adult flies develop grossly mishapen scutellums with severe macrochaete patterning defects. Three representative thoraxes are shown.

To test the hypothesis that the *nab* DWME is licensed in part by Ap to respond to Notch and BMP developmental signals only in the dorsal wing compartment, we crossed our *nab* DWME reporter line into backgrounds differing in the dosage of normal Ap. We observed that DWME-driven GFP reporter expression collapses to two stellate clusters of just a few cells along the D/V border region as the levels of normal Ap is reduced. (Fig. 6B–D). This result is more consistent with Ap being required to allow maintenance of DWME activity in off-margin descendants of a few margin cells. However, further studies will be needed to explore this, the order of binding events, and the effects on nucleosome positioning.

### Dorsal-compartment expression of nab is required for normal patterning of wing and thorax

Expression of *nab* in the dorsal compartment of the developing wing pouch is the result of cells using DWME and/or QE activity. To understand better the role of the DWME versus the QE, we used the *ap-GAL4* hypomorph (*ap^GAL^*^4^) to drive *nab* RNAi. This likely brings down *nab* mRNA levels in the dorsal compartment while also specifically disrupting the Ap-dependent activity of the endogenous *nab* DWME. Thus, in this experiment the most severe disruption of *nab* expression is expected to be in the DWME-active cells.

Using one copy of the *ap^GAL^*^4^ allele to drive expression of UAS:*nab*-RNAi, we observe a consistent set of phenotypes that include an exaggerated creased wing along the vein II (Fig. 6D). This is often accompanied by a malformed delta-like junction between the first and second wing veins at the anterior margin (Fig. 6E). Sometimes this is accompanied by a disrupted bristle pattern at the juncture of the margin (Fig. 6E vs. 6F wild-type). In addition, the scutellum of these flies is malformed and diminished in size with typically one or more anterior and posterior scutellar bristles missing (compare Fig. 6G,H to Fig. 6I—K). In these flies, the anterior and posterior dorsocentrals in the notum are also frequently absent (Fig. 6I—K). Thus, there are missing or mis-specified macrochaetes in both the notum and scutellum. Wings from *ap^GAL^*^4^:*nab*-RNAi adults show other defects including occasional ectopic wing veins, gaps in the wing vein patterning, and diminished or absent anterior cross-veins (data not shown). Therefore, we propose that normal patterns and/or levels of *nab* are required in the dorsal margin cells in order to ensure proper wing vein specification and patterning.

The results of dorsal compartment-specific knockdown of *nab* suggests that expression levels are critical to morphogenetic pattering of the notum and wing veins (Fig. 6). To explore further the importance of *nab* expression in wing imaginal discs, we expressed *nab*-RNAi in two additional ways. First, we used an endogenous *GAL4-nab* enhancer trap line (nab-NP3537) to drive the same *nab*-RNAi transgene. The prediction is that in this experiment, both endogenous Nab and Gal4 proteins are being synthesized simultaneously as the result of the different regulatory enhancers at Nab inducing transcription of both *nab* and *GAL4*. Thus, this experiment might be expected to result in only a partial knockdown of *nab* protein because of the delayed transcriptional/translational cycle of Gal4-driven *nab*-RNAi. In this experiment, we observe the mildest effect of all the *nab*-RNAi experiments in that we only observed one phenotype: frequent loss of the posterior cross-veins but only in adult females (see arrows in Fig. 7 B and C).

**Figure 7.**
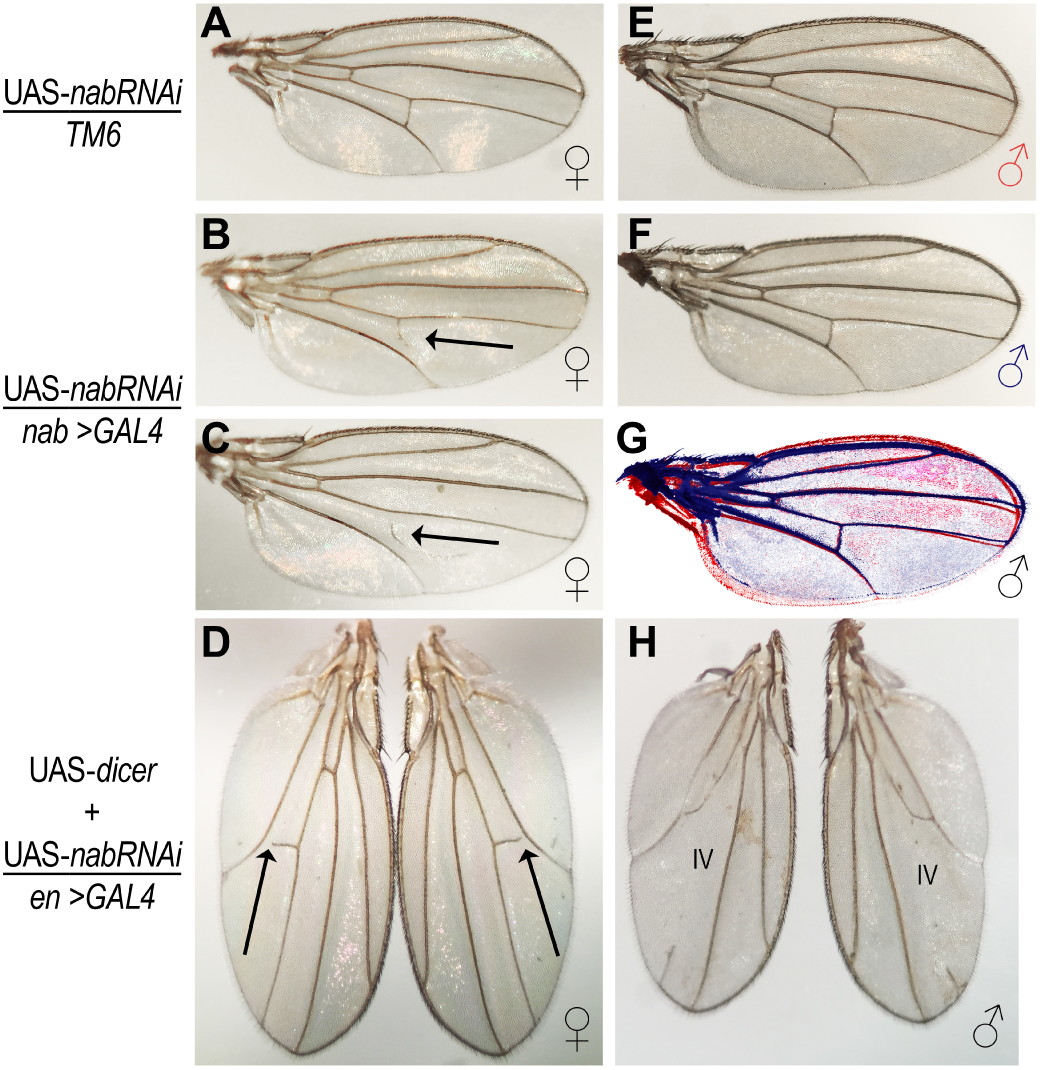
Distinct levels of *nab* expression are required for normal developmental patterning of the notum, scutellum, and wing veins. Wing vein patterning in female **(A–D)** and male **(E–H)** fly wings carrying a UAS-*nab*-RNAi transgene without a GAL4 driver (A, E, and the red wing outline in G), a *nab* locus GAL4 enhancer trap driver (B, C, F, and blue wing outline in G), or an *en* locus GAL4 enhancer trap driver augmented with additional UAS-*dicer* expression (**D, H**). Posterior cross-vein are often lost or incomplete in female wings with *nab*-RNAi knockdown arrows in **(B–D)**. The fourth wing vein is often lost in male wings **(H)**. Thus, normal patterns of *nab* expression in both the dorsal and ventral compartments are crucial to wing vein patterning.

To test the role of *nab* expression in other wing disc compartments, we used an *engrailed* (*en*) *GAL4* enhancer trap line to drive *nab*-RNAi in the posterior compartment, which spans both D/V compartments (see Material and Methods). En is a selector protein expressed early in the posterior compartments of wing imaginal discs, where it determines posterior-compartment identity (growth/size, shape, and wing vein patterning) (Lawrence and Morata 1976; Kornberg 1981; Blair 1992; Brower 1986). Importantly, in the *en-GAL4* experiment, *nab*-RNAi would be induced and maintained prior to transcription of *nab* in early third instar wing imaginal discs (Terriente Félix *et al*. 2007). To increase the severity of any phenotypes, we used the *en-GAL4* transgene to drive both *UAS-nab*-RNAi and *UAS-dicer*. This experiment results in a more severe wing vein patterning defect in terms of frequency in both females and males (Fig. 7 D and H, respectively). This phenotype, which includes incomplete and missing longitudinal veins as well as cross-veins, is restricted to the posterior compartment. We do not see any morphological defects in scutellar development, suggesting that *ap* hypomorphs sensitize scutellar development to loss of Nab function.

In summary, *nab*-RNAi knockdown experiments demonstrate that *nab* expression is important in all compartments of wing imaginal discs, including both proximal and distal parts of discs, for the normal developmental patterning of the scutellum, thoracic macrochaetes, and wing veins. This is consistent with *nab*’s intronic complex of three wing imaginal disc enhancers (QE, PWE, and DWME).

### The nab DWME is a lineage specific enhancer maintaining expression in off-margin clonal descendants of margin cells

To better understand the individual roles of the *nab-A* binding sites mediating Notch and Dpp signaling, we first examined the developmental onset of expression in wing imaginal discs. We find that DWME-driven expression begins around late second/early third instar discs in a few cells at the margin (Fig. 8 A and B). This is consistent with the earliest reported detection of Nab protein in early third instar discs (Terriente Félix *et al*. 2007). As the discs develop, there are many examples of dividing daughter cells in which one of the daughter cells is dividing away from the D/V boundary while still maintaining equivalent amounts of expression (see circled pairs of cells in Fig. 8A′–D′, and I′). These pairs of dividing cells can also be seen in later stages when descendants are at much farther distances from the margin (Fig. 8I′). We also observe decreasing DWME-driven reporter activity in early to mid third instar cells along the margin when peak p-Mad is presumably still centralized as a single (*i.e*., non-bimodal) peak (see series of arrows of decreasing size in (Fig. 8I′).

**Figure 8.**
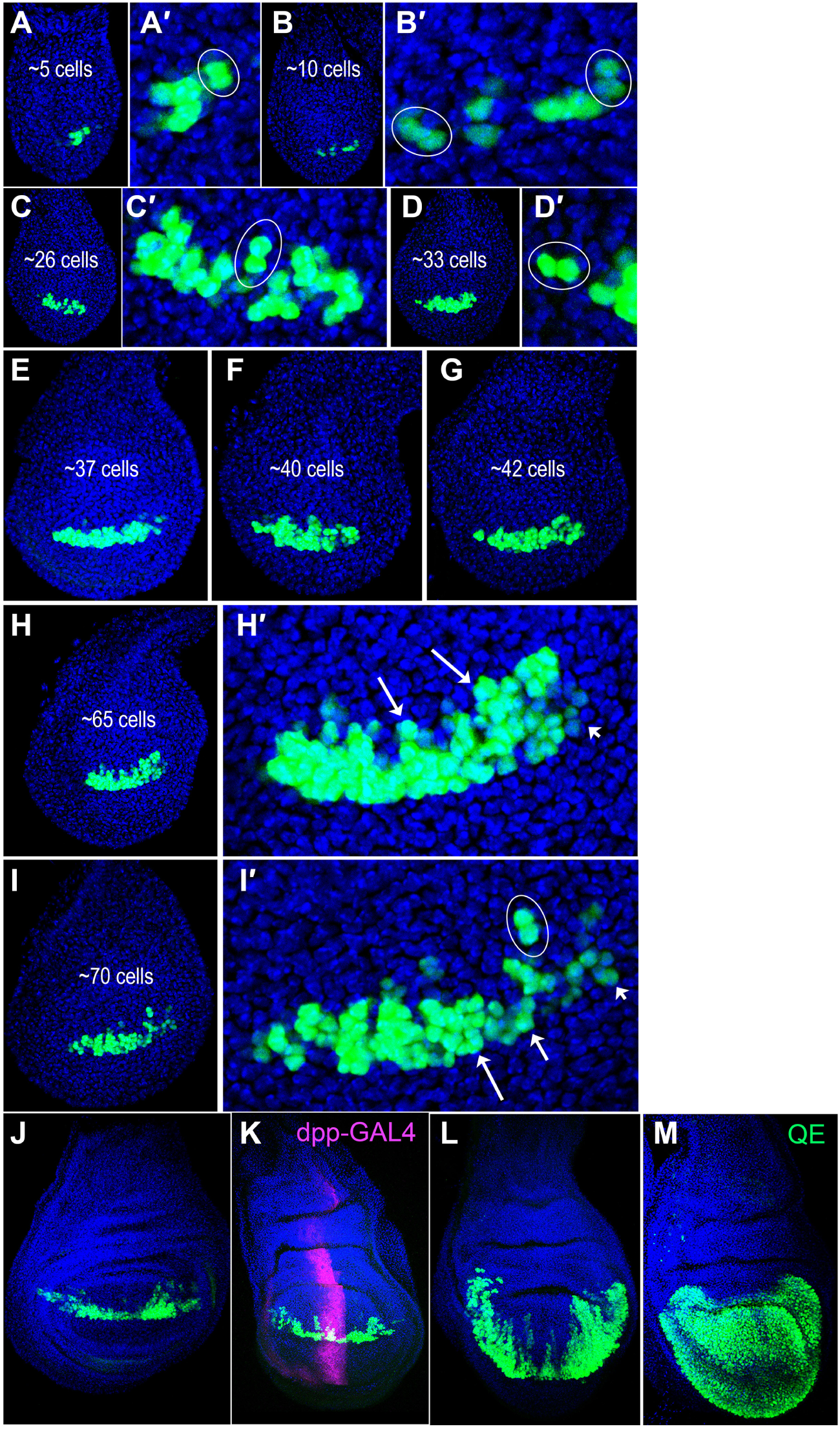
The *nab* DWME is stably maintained in off-margin clonal descendants. **(A–L)** GFP reporter activity driven by the *nab* dorsal wing margin enhancer (DWME, *nab-A* fragment) starting during late second instar to late third instar wing imaginal discs are shown arranged according to the number of expressing cells based on nuclear GFP intensity. **(A**′–D′,H′,I′) Zoomed-in images. Inspection of recently divided daughter cells (*e.g*., ovals in A′,B′,C′,D′, and I′) demonstrate that GFP activity is equally maintained in both cells even when the direction of cytokinesis places one of the daughter cells farther away from the dorsal compartment boundary. Cells far removed from the D/V boundary but still near the A/P compartment boundary display robust GFP expression (large arrows in H′). However, cells further away from the A/P compartment boundary display decreasing levels of GFP expression even when they are on the dorsal margin itself (smaller arrows in H′ and I′). During early-late third instar **(J,K)** and late-late third instar/pre-pupal **(L)** discs, the DWME activity is maintained in clonal descendants of dorsal margin cells that are now located deep into the wing pouch. We thus propose that the DWME functions is a lineage-specific margin enhancer that maintains expression in dorsal off-margin clonal descendants. The disc in **(K)** is doubled-stained for Dpp (magenta). **(M)** The *nab* quadrant enhancer (QE, *nab-B* fragment) is noticeably weaker or not active in dorsal compartment off margin cells from late third instar/pre-pupal discs. Representative disc shown for comparison.

This temporal profile of *nab* DWME activity suggested to us that the DWME marks a clonal patch of cells descended from a margin cell, which experienced early Notch-Delta signaling (see Fig. 9B and Discussion). This would also explain a role for the Dpp input in stabilizing activity of the DWME in clonal descendants removed from the border of active Notch-Delta signaling. Like clonal fate mapping in wing imaginal discs (Dahmann and Basler 1999), cells expressing DWME-driven GFP are patch-like clusters of dividing cells that respect both D/V and A/P compartment boundaries. Consistent with this idea, DWME-driven patches of GFP expression resemble the D/V-oriented cell proliferation patterns made by marked clones in wing imaginal discs (González-Gaitán *et al*. 1994). To explore this idea further, we also looked at the *nab* QE (*nab-C*) reporter in late third instar/pre-pupal wing imaginal discs specifically to see if the deficiency in dorsal margin expression becomes more noticeable over time. We find that this is possibly the case as the margin gap appears more pronounced than earlier stages (Fig. 8M). This suggests that the *nab* DWME and QE are complementary lineage-specific enhancers that have evolved to deliver Nab expression in proliferating wing pouch cells more uniformly than what the QE alone could drive.

**Figure 9.**
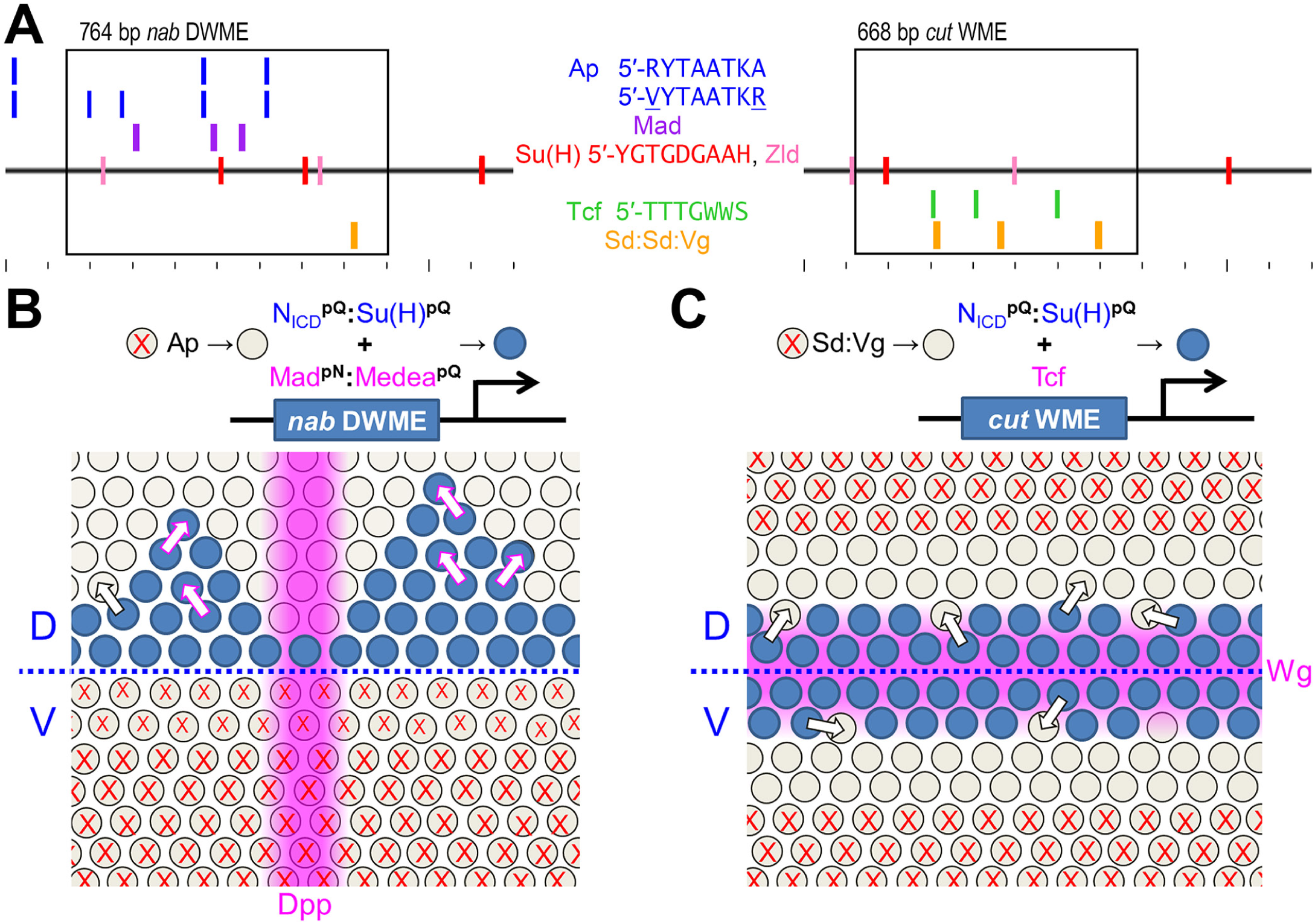
An enhancer model featuring selector licensing for pQ-mediated-signal integration. **(A)** Shown are the trascription factor binding site motif distributions for the *nab* dorsal wing margin enhancer (DWME, left) and the *cut* wing margin enhancer (WME, right), both of which are induced by Notch signaling and Su(H) binding sites. As much as possible, matches to the indicated motifs are shown on separate tracks for ease of visualization. Tick marks represent 100 bp intervals. Boxes represent the minimalized enhancers, but in both cases augmented expression is seen with larger fragments. **(B, C)** Below each enhancer are models of how each enhancer works in the context of both homeotic licensing and graded pQ signal integration or lack thereof. In both examples, selectors are envisioned as allowing certain cells (cell without X’s) to be receptive to transcriptional effectors of signaling pathways (Notch, Dpp, and Wg). The pQ/pN-rich Mad:Medea complex are envisioned as stabilizing the active DWME in daughter cells of margin cells (B). In contrast, no such stabilizing effect is envisioned for the *cut* WME, which lacks both Mad:Medea sites (purple boxes) and instead has only Tcf (green boxes) and Su(H) (red boxes) binding sites (C). Thus, this enhancer would become inactive in margin daughter cells that divide away from the margin border, where there is active Notch-Delta signaling.

## Discussion

We used a comparative genomics approach to identify and characterize a novel enhancer mediating tissue-compartment specific activation based on the integration of two signaling pathways: Notch and BMP/Dpp. This approach led us to a dense intronic cluster of binding sites for the desired activators and selectors at the *Drosophila* gene *nab*. “Nab” was named for “NGFI-A binding” and is present in vertebrates as Nab1 and Nab2, where they work as co-repressors of Egr-1 (NGFI-A) and Krox20 to control proliferation, patterning, and differentiation (Russo *et al*. 1995; Svaren *et al*. 1996). Similarly in the nematode C*. elegans*, this gene encodes MAB-10/NAB, which is a co-repressor of LIN29/EGR and is involved in cellular differentiation (Harris and Horvitz 2011). In *Drosophila*, LIN-29/EGR functions appear to be maintained across three paralogous genes *rotund* (*rn*), *squeeze*, and *lin19* (Vilella *et al*. 2009). Correspondingly, *Drosophila* Nab functions as a co-repressor of Rn in wing imaginal discs and as a co-activator of Squeeze in a subset of embryonic neuroblasts (Terriente Félix *et al*. 2007). Thus, *nab* is conserved across bilaterian animals and encodes a co-factor of a subfamily of C2H2 zinc finger transcription factors involved in developmental patterning and differentiation.

The dense site cluster that we identified in the first intron of *nab* drives a novel expression pattern that is consistent with integration of a D/V Notch signal and an orthogonal A–P BMP signal. The *nab* locus is already known to be induced by Dpp/BMP signaling (Ziv *et al*. 2009; Hadar *et al*. 2012). This is consistent with our finding several canonical Mad:Medea binding sequences in the *nab* DWME (Fig. 3A) and several low-affinity Mad:Medea binding sequences in the *nab* QE (not shown). Furthermore, both the DWME and QE drive A–P modulated expression in the wing pouch (Fig. 4B, C, B′″, and C′″). Here, we investigated the role of Notch signaling as one input integrated by the *nab* DWME. We find that the *nab* DWME has at least two canonical Su(H) binding sites that are critical to its activity at the D/V margin. This activity is also impacted by a mutant *Notch* background. Thus, this may be the first example of a wing disc enhancer that is downstream of both Notch and BMP signaling. Furthermore, by virtue of its obligate Notch signaling input, the DWME’s transcriptional readout of the BMP morphogen gradient is an idealized two-dimensional graph of p-Mad levels (*y*-axis) at different A–P positions (*x*-axis).

To deconstruct the developmental role of *nab* expression in different wing compartments, we used the *ap-GAL4* enhancer trap to drive *nab*-RNAi in the dorsal compartment. This RNAi experiment would also knockdown activity of the Ap-dependent dorsal wing margin enhancer in the dorsal wing compartment. With this manipulation, we observed morphogenetic defects of both the thorax and the wing, suggesting an important role for Nab in developmental patterning of the dorsal compartment. More specifically, the patterning defects affect the scutellum and thoracic macrochaetes, consistent with both *nab* expression and its proximal wing imaginal disc enhancer (PWE). This experiment also resulted in an increased rate of wing vein patterning defects. We also observed similar wing vein patterning defects, including loss of the posterior cross-vein in males and females and loss of longitudinal veins IV and V in males, when we drove *UAS:nab*-RNAi in the posterior compartment using an *en-GAL4* enhancer trap line. A milder knockdown using a *nab-GAL4* enhancer trap line resulted in an increased loss of the posterior cross-vein in adult females. Thus, of all the phenotypes, developmental specification of the posterior cross-vein was the most sensitive to diminished Nab function. This is a significant finding given that *crossveinless c* was the only other locus in the genome having a similar cluster of binding sites as the *nab* DWME (Table 1).

### Lineage-specific aspects of wing margin and wing quadrant enhancers

Inspection of the site composition of the *nab* DWME shows it is a lot like the wing margin enhancer (WME) at *cut* (Fig. 9A). Both wing margin enhancers are induced by Notch signaling and have functional sites for Su(H) (Jack *et al*. 1991; Neumann and Cohen 1996a). We find both enhancers also have sequences matching the binding preferences for Zelda and Scalloped dimer dimers sites, although the *cut* WME has many more of the latter. However, the *nab* and *cut* wing margin enhancers each have multiple binding sites either for Mad:Medea or for Tcf, but not for both (compare purple SMAD sites in the *nab* DWME versus green Tcf sites in the *cut* WME in Fig. 9A). Unlike the Mad:Medea complex, *Drosophila* Tcf/Pangolin and its *β*-catenin co-activator Armadillo, which together mediate Wg/WNT signaling, are not pQ/pN-rich. We thus propose the following model for pQ-mediated Notch/BMP signal integration at the *nab* DWME (Fig. 9B) that distinguishes it from the *cut* WME (Fig. 9C).

We hypothesize that pQ/pN-mediated aggregation of Mad:Medea and Su(H):NICD complexes produced by the *nab* DWME stabilizes and maintains an active DWME in off-margin clonal descendants of D/V margin cells. Thus, the *nab* DWME would remain active only in daughter cells that are also simultaneously experiencing high levels of nuclear p-Mad (see arrows in Fig. 9B). In contrast, the *cut* WME would become inactive in these same off-margin daughter cells because they are no longer experiencing Notch or Wg signaling (see arrows in Fig. 9C). In this way, the *nab* DWME functions as a lineage-specific enhancer in regions with active phosphorylated-Mad (p-Mad). In contrast, the *cut* WME is not lineage-specific because its activity is extinguished in cell descendants as they divide away from the margin. Consistent with this model of Notch/BMP integration at the *nab* DWME, the presumed p-Mad input is not sufficient for driving DWME expression in cells that although having peak p-Mad levels are not descended from margin cells.

The *cut* locus is not expressed throughout the wing pouch like the *nab* and *vg* genes, both of which have separate margin/boundary and quadrant enhancers. This also highlights the lineage-specificity of some wing pouch enhancers as follows. The *vg* locus contains a Notch-target boundary enhancer (Williams *et al*. 1994) as well as a separate BMP-target quadrant enhancer (Kim *et al*. 1997). The *vg* boundary enhancer drives expression broadly throughout the presumptive wing border region while the *nab* DWME drives expression in a dorsal margin cells and their off-margin descendants. While the *vg* locus has separate “single-channel” enhancers responsive to either Notch or BMP signaling pathways, the *nab* locus has enhancers responsive to combined Notch/BMP signals and (presumably) BMP-only signals. Nonetheless, separate margin/boundary and quadrant enhancers would be required by loci expressed uniformly in proliferating cells of the developing wing pouch. Potentially, this “rule” might generalize across diverse developmental systems in which a gene is expressed broadly in a proliferating tissue patterned by a smaller number of compartment organizer cells. One enhancer would respond to signals experienced specifically by organizer cells, and a second enhancer would respond to signals experienced more broadly across the developmental field.

Our study of the *nab* intronic enhancer complex also demonstrates a remarkable aspect of transcriptional enhancers. We found that all of the *nab* intronic enhancers are also silencers. We discovered this because we were monitoring expression across several tissues of the embryo, larva, and pupa. This suggests that we should be cautious about the effects of enhancer (“over”)minimalization. This hidden aspect of enhancers may predispose tissue-specific enhancer screens to high false-positive rates of detection if short regulatory DNA fragments are associated with ectopic activities. In any case, these results suggest a new role for Su(H) in mediating inter-enhancer silencing and attenuation. This functional role is worth testing in greater depth across several additional examples.

### Selectors: Licensing Factors via Nucleosomal Positioning?

Homeotic selectors were defined initially as TFs that decide and maintain a choice of alternate cell fate potentials for a clonal population of cells and their descendants (García-Bellido 1975; Lawrence *et al*. 1979). One early clue suggesting that they correspond to a specific regulatory mechanism was that (homeotic) selectors in animals were predominantly found in the form of homeodomain-containing factors (McGinnis *et al*. 1984). While selectors were initially discussed in terms of cell fate determination and cell fate potential, a consideration of evolutionary processes suggested an emphasis on the role of transcriptional enhancers as important components of selector-based regulation (Akam 1998). Thus, we hypothesize that the phenomenology of selectors acting on cell fate decisions lies in their roles as factors licensing enhancer DNAs to receive graded developmental signals through any number of pathways (*e.g*., Notch, BMP/Dpp, WNT/Wg, and Hh).

The *nab* DWME, *nab* QE, *cut* WME, and *vg* WME (*i.e*., the boundary enhancer, BE) have binding sites for homeotic selectors in addition to those for the pathway-mediating effectors. Selectors can act in a negative or positive manner as demonstrated by known wing disc enhancers. The *vg* WME uniquely has matching sites for the posterior Hox factor Abd-B homeotic selector (5′-YYTTTATGK) that are not found in either the *nab* or *cut* WMEs (data not shown). Abd-B plays a negative selector role by prohibiting expression of *vg* in posterior segments (Carroll *et al*. 1995). Similarly, the *nab* QE possesses binding sites for Homothorax (Hth) (5′-ARYDATSRC), which is known to ChIP to this enhancer (Slattery *et al*. 2013). Hth is expressed throughout the disc except in the wing pouch and is thus likely to prohibit expression outside of the wing pouch. This would be consistent with the clean border of *nab* QE reporter expression at the wing pouch/hinge border. In contrast, Hth sites are absent in the *vg* WME, which drives D/V margin expression beyond the pouch.

The wing margin enhancers at *nab* and *cut* are likely to use positively acting selectors. Consistent with the site composition of the *nab* and *cut* WMEs (Fig. 9A), the former would be positively licensed primarily by the selector Ap for the dorsal wing compartment (see cells without X’s in Fig. 9B), while the latter would be positively licensed by the selector complex of Sd and Vg (see cells without X’s in Fig. 9C). Thus, a reduction of the dosage of Ap causes DWME-driven expression to collapse to a few cells along the wing margin. Because of the dorsal compartment-specific expression of Ap, and the known localization of activity of Notch signaling to the margin, it is the Notch pathway, secondarily augmented by the BMP input, that drives its expression pattern. Ap only marks a larger domain in which DWME activity can occur. Thus, we interpret this as Ap “licensing” the Notch-dependent DWME for expression exclusively in the dorsal compartment. Additional positively-acting selectors are likely to work at the *nab* DWME based on conservation of known binding sequences, but further work will be required to elucidate these more definitively. In addition to the cluster of Ap binding sites, and a single Sd:Sd:Vg site, we also find a binding site for the homeotic selector Dll (5′-ATAATYAT), which has a similar expression pattern to Sd and Vg at the wing margin (Campbell and Tomlinson 1998), but this site is not as distinct as the Sd dimer sites. We thus suspect that the DWME might be licensed collectively by Sd, Vg, Dll, and Ap.

We speculate on how the selector licensing class might function to permit and restrict enhancer activities based on certain sequence features in the *nab* DWME. These features hint at the role of selectors in making the enhancer sequences either receptive or refractory to effector DNA binding via fixed or remodeled nucleosomal positions. The core sequence of the DWME possesses contains fixed nucleosomal positions in early embryonic stages prior to DWME activity (Fig. S4 in Supporting File S3) (Mavrich *et al*. 2008; Langley *et al*. 2012). At specific sites flanking these fixed nucleosomes, there are also polyT, homopolymeric runs (Fig. S4), which can act as nucleosome repulsing sequences (Anderson and Widom 2001; Segal and Widom 2009). In addition, we also see evidence for competing TT/AA/TA dinucleotide phasing embedded at these fixed nucleosomal windows (Lowary and Widom 1998; Thåström *et al*. 1999; Segal *et al*. 2006). Based on the known Su(H) binding structures (Arnett *et al*. 2010), these fixed positions would make the central S1 Su(H) site unavailable for binding (Fig. S4). We thus suggest that some selectors may act by simply binding and stabilizing alternate nucleosomal positions that reveal effector binding sites. Intriguingly, we also find that Apterous binding sites coincide with the nucleosomal DNA entry regions that are also associated with the repressor linker histone H1 (Syed *et al*. 2010). Consistent with the inability of Su(H) to bind the DWME sites in the absence of Ap, we find that Su(H) ChIP signals at the *nab* DWME have not been detected in embryonic stages (Nègre *et al*. 2011).

We speculate that the above homeodomain-containing selectors (Ap, Dll, Hth) and other selectors (*e.g*., Sd and Vg) work via a mechanism that is different from the graded spatial (Notch/BMP) and temporal (Zld) patterning effectors. The above set of selectors are deficient in pQ (Fig. 1A), pN (Fig. 1B), and mixed p(Q/N) carboxamide (Fig. 1C) tracts unlike the spatiotemporal effectors and the Mediator co-activator complex, which they recruit in their role as activators. We suggest that the phenomenon of graded signal integration by effectors also occurs via generic poly-carboxamide aggregation as dictated by the composition of binding sites present in different DNA enhancer scaffolds. In stark contrast, pQ/pN-deficient selectors could bind and fix (*i.e*., stabilize) default or alternate nucleosomal positions without cross-interacting with pQ/pN-rich effectors. Furthermore, the relationship between selector and pathway effector binding sites would be informed by a helically-phased “regulatory reading frame” based on nucleosomal positioning.

## Acknowledgments

This work was supported in part by an NSF CAREER award to AE to study morphogen gradient enhancer readouts (IOS:1239673), an NIH T-32 bioinformatics training grant support to ES, and two Evelyn Hart Watson research fellowships (2014 and 2015) to ES. We thank Timothy Fuqua for assistance with some of the antibody staining protocols. We thank Timothy Fuqua for comments on an earlier version of this manuscript.

